# Programmed electrical stimulation in human iPSC-derived cardiomyocytes reveals mechanisms of lethal arrhythmias in Calcium Release Deficiency Syndrome

**DOI:** 10.64898/2026.04.09.717574

**Authors:** Saif F. Dababneh, Alia Arslanova, Mariam Butt, Torin Halvorson, Thomas M. Roston, Jason D. Roberts, Seiko Ohno, Farah Jayousi, Philipp F. Lange, Leif Hove-Madsen, Robert A. Rose, Edwin D.W. Moore, Filip Van Petegem, Shubhayan Sanatani, S.R. Wayne Chen, Glen F. Tibbits, Maksymilian Prondzynski

**Author notes:** Author deceased (Glen F. Tibbits).

## Abstract

**Background:** Calcium release deficiency syndrome (CRDS) is a recently described inherited channelopathy caused by loss-of-function variants in *RYR2*. Clinically, CRDS patients present with lethal ventricular arrhythmias which are not reproduced on exercise stress testing, unlike catecholaminergic polymorphic ventricular tachycardia. A hallmark trigger identified for CRDS mimics a long-burst, long-pause, short-coupled extra-stimulus (LBLPS) programmed electrical stimulation protocol, which was experimentally validated in humans and mouse models. Moreover, application of a long-burst, long-pause (LBLP) protocol alone can induce an abnormal repolarization on the first sinus beat that is unique to CRDS. However, the electrophysiological basis of CRDS in human cardiac tissue, including other triggers, are not fully understood, and whether clinically relevant arrhythmias can be observed in human stem cell models remains unknown.

**Methods:** We performed electrophysiological and arrhythmia inducibility studies using clinically relevant programmed electrical stimulation protocols in two-dimensional cardiac tissue generated from metabolically matured human induced pluripotent stem cell-derived cardiomyocytes (hiPSC-CMs) carrying the CRDS variant RyR2-E4146D. High spatiotemporal optical mapping and multielectrode arrays were used for electrophysiological phenotyping.

**Results:** At baseline, E4146D^+/-^ monolayers showed no arrhythmias, similar to controls. During rapid pacing, E4146D^+/-^ promoted electrical vulnerability by reducing the threshold for action potential duration (APD) alternans and Ca^2+^ alternans and increasing the propensity for spatial discordance of alternans. In response to LBLP pacing, E4146D^+/-^ monolayers often demonstrated an abnormal repolarization response characterized by spatially dispersed APD prolongation and large Ca^2+^ release. Notably, LBLPS pacing produced early-after depolarization (EAD)-driven triggered activity resulting in re-entrant tissue conduction patterns, explaining the short-coupled ectopy driven arrhythmias seen in CRDS patients. Similar arrhythmias were observed when EADs developed during spatially discordant alternans. Lastly, flecainide showed efficacy in suppressing arrhythmia inducibility for the here studied variant.

**Conclusions:** We developed the first hiPSC model for CRDS which recapitulates clinically observed and inducible arrhythmias. Our model provides novel insights into tissue-level, re-entrant arrhythmias, which are initiated by EADs during electrically vulnerable states in CRDS human cardiac tissue and can be suppressed by flecainide. This model provides the framework for studying other CRDS variants and complex arrhythmias in hiPSC-CMs and establishes a human-based new approach method (NAM) for drug and gene therapy development for CRDS.

**CLINICAL PERSPECTIVE:** *What is new?:*

▪ We developed the first human stem cell-derived cardiomyocyte (hiPSC-CM) tissue model for calcium release deficiency syndrome (CRDS) which recapitulates its hallmark clinical features, including inducible ventricular arrhythmias with programmed electrical stimulation and post-pacing repolarization abnormalities.
▪ Using genome edited and metabolically matured hiPSC-CMs combined with high spatiotemporal optical mapping, we show that tissue-level arrhythmias are initiated by early-after depolarizations (EADs) which develop during electrically vulnerable states, leading to re-entrant conduction patterns. We comprehensively characterize the features of EAD-induced triggered activity, showing that these ectopic beats promote re-entry through slower conduction velocities and shorter action potential durations. This uncovers how EAD-induced short-coupled ectopy leads to malignant ventricular arrhythmias in CRDS patients, and establishes the phenotype for future hiPSC-CM investigations.
▪ We identified flecainide as an effective agent in suppressing arrhythmias on single cell and tissue levels in hiPSC-CMs for this CRDS variant, reproducing clinical results.

*What are the clinical implications?:*

▪ CRDS has only recently been described as a unique channelopathy caused by loss-of-function *RYR2* variants, and much of its triggers and mechanisms in human cardiomyocytes remain unclear. The arrhythmias observed are often not related to exercise, and exercise stress testing does not reproduce these abnormalities. No human models exist to date which closely recapitulate the triggers shown to induce tissue-level arrhythmias in patients and mouse models. Our model demonstrates that programmed electrical stimulation, without pharmacological β-adrenergic stimulation, can reliably induce the same arrhythmias seen clinically, enabling accurate disease modeling and drug development.
▪ Combining programmed electrical stimulation in cardiac tissue derived from genome-edited hiPSC-CMs with high spatiotemporal optical mapping is a robust and novel approach to identify the mechanisms of complex, tissue-level arrhythmias which remain underexplored, such as short-coupled ventricular fibrillation, in a patient-specific and translational manner.

## INTRODUCTION

Calcium release deficiency syndrome (CRDS) is a recently described inherited arrhythmia syndrome caused by loss-of-function (LOF) variants in the cardiac ryanodine receptor (RyR2).^1^ RyR2 is a central regulator of excitation-contraction coupling linking membrane depolarization to sarcoplasmic reticulum (SR) Ca²⁺ release. Clinically, CRDS patients present with ventricular arrhythmias or sudden cardiac arrest typically without structural heart disease.^2^ Unlike patients with catecholaminergic polymorphic ventricular tachycardia (CPVT), an inherited channelopathy primarily caused by gain-of-function *RYR2* variants,^3^ CRDS patients often lack adrenergically induced arrhythmias and appear normal on exercise stress testing, rendering this lethal disorder clinically concealed using standard diagnostic testing.^2,4^ Given that CRDS is in its infancy, its prevalence remains unknown. Prior to the recognition of CRDS as a disease entity, LOF *RYR2* variants have been linked to several phenotypes including long QT syndrome (LQTS),^5^ short-coupled ventricular fibrillation (scVF) or torsades de pointes (scTdP),^5,6^ and idiopathic ventricular fibrillation (IVF).^7,8^ Additionally, CRDS patients are at risk of mis-and underdiagnosis due to phenotypic overlap with CPVT, with around 13% of CPVT probands presenting with unexplained cardiac arrest without any inducible arrhythmia at exercise stress tests, suggesting potential CRDS misdiagnosis.^9^

In CPVT, leaky RyR2 channels result in diastolic SR Ca²⁺ leak driven by adrenergic stress, leading to NCX-mediated delayed afterdepolarizations (DADs) which clinically progress from premature ventricular contractions, bigeminy, to polymorphic ventricular tachycardia.^10,11^ Functional studies in heterologous systems and isolated mouse CMs demonstrated that CRDS-linked variants reduce RyR2 channel activity.^1,12,13^ A D4646A^+/-^CRDS mouse model recapitulated attenuated RyR2 function and confirmed the absence of catecholamine-induced arrhythmias, mirroring the clinical phenotype. Instead, ventricular arrhythmias in CRDS were unmasked by programmed electrical stimulation using the long-burst, long-pause, and short-coupled extra-stimulus (LBLPS) protocol, which has been validated in mice and CRDS patients.^1^ Based on mechanistic mouse studies, CRDS is thought to be driven by early after depolarizations (EADs) which result in re-entrant arrhythmias.^1,14^ More recently, using only the long-burst portion (i.e., rapid pacing) in patients and mice was found to result in hallmark repolarization abnormalities unique to CRDS compared to patients with either CPVT, supraventricular tachycardia, or those with unexplained cardiac arrest.^15^ This is a promising diagnostic test that is currently being explored in a multi-centre international study (DIAGNOSE-CRDS, NCT06188689).

One model proposes that RyR2 LOF elevates the threshold for store overload-induced Ca²⁺ release (SOICR). This process is driven by excessive SR Ca²⁺ accumulation, due to a reduced physiological Ca²⁺ leak during rapid pacing, until abrupt and large depolarization-induced Ca²⁺ release occurs, triggering ventricular arrhythmias through NCX-mediated EADs and re-entrant activity.^1^ However, the cellular mechanisms in human cardiac tissue remain unclear.

The use of human induced pluripotent stem cell-derived cardiomyocytes (hiPSC-CMs) has shown great promise in modeling inherited arrhythmias in a personalized and scalable fashion^16,17^ with several studies demonstrating feasibility and impact for conditions such as Long QT Syndrome (LQTS),^18,19^ Brugada syndrome,^20^ NAA10-deficency syndrome,^21^ and CPVT.^22–24^ Especially, RyR2-CPVT hiPSC-CM models were able to recapitulate adrenergic stimulation-mediated DADs enabling drug testing efforts in a patient-specific manner.^22,25,26^ Moreover, novel gene therapy approaches for RyR2-CPVT have been developed and validated in hiPSC-CM models.^22,27,28^ However, modeling complex arrhythmias *in vitro* remains challenging and the majority of studies do not focus on tissue-level arrhythmia mechanisms. A few studies have previously investigated LOF *RYR2* variants in single hiPSC-CMs, showing arrhythmic Ca^2+^ transients with and without β-adrenergic stimulation.^29–32^ However, it is unclear how well these phenotypes mimic the patients’ clinical presentations. In some of these studies, β-adrenergic stimulation increased the propensity for DADs,^30–32^ which is inconsistent with mouse studies showing an increased propensity for EADs in CRDS cardiomyocytes.^1,14^ It remains unclear whether the inconsistency is due to differences in species, immaturity of hiPSC-CM Ca^2+^ handling, or other factors. Additionally, previous hiPSC-CM studies have not tested arrhythmia inducibility using programmed electrical stimulation protocols such as LBLPS, which is clinically used for inducibility and diagnosis of CRDS patients.^1,33^

Here, we demonstrate the utility of clinically validated programmed electrical stimulation protocols to induce arrhythmias in monolayers of metabolically matured hiPSC-CMs and provide novel mechanistic insights into CRDS pathophysiology. We utilized CRISPR-Cas9 genome editing to generate an hiPSC line carrying a previously reported LOF *RYR2* variant, E4146D^+/-^,^5^ which was linked to LQTS and scVF/scTdP. We used high spatiotemporal resolution optical mapping to measure voltage and Ca²⁺ dynamics, revealing reproducible induction of arrhythmias in our model. By applying programmed electrical stimulation protocols in hiPSC-CM tissue, we show how re-entrant arrhythmias are triggered and propagated in human CRDS tissue. Our study defines the electrophysiological mechanisms underlying inducible arrhythmias in CRDS in human cardiomyocytes and supports the potential efficacy of flecainide as a treatment for CRDS.

## METHODS

### Ethics

The clinical study protocol was approved by Shiga University of Medical Science (R2011-128). Written informed consent was obtained from the patient and parents. Biosafety ethics approval was obtained from the University of British Columbia (B25-0223) for all *in vitro* studies.

### Genome editing in human pluripotent stem cells

The E4146D variant in *RYR2* was introduced into low-passage (<20), commercially obtained hiPSCs (WiCell, STAN248i-6717C1) using Cas9 RNP electroporation. STAN248i-6717C1 is derived from a healthy female participant who was 38 years old at time of collection, and the cell line has undergone rigorous quality control (documentation found at provider’s webpage). Cas9-RFP was complexed with a custom designed sgRNA with a 98/100 off-target score (Benchling) at equimolar concentrations to form RNP complexes (Supplementary Table 1). A single-stranded oligodeoxynucleotide (ssODN) donor template was included to direct homology-mediated repair sequence (Supplementary Table 1). hiPSCs were dissociated into single cells using Accutase (STEMCELL Technologies, 07920) and resuspended in Opti-MEM Reduced Serum Medium (Gibco, 31985062) for electroporation. Following electroporation in a 0.1 cm gap cuvette (Bio-Rad, 1652083), cells were plated into mTeSR Plus (STEMCELL Technologies, 100-0276) supplemented with 10 µM ROCK inhibitor (Y-27632, Biogems, 1293823) for recovery. Sixteen hours post-electroporation, RFP-positive cells were isolated by FACS using the BD FACSAria II and plated into 6-well plates preconditioned with CloneR supplement (STEMCELL Technologies, 05888) and penicillin-streptomycin (Gibco, 15140148) to enhance single-cell survival. Colonies were manually picked 4-6 days post-FACS and expanded until confluent. One plate of each clone was used for DNA extraction and genotyping, while a replicate plate was cryopreserved in mFreSR (STEMCELL Technologies, 05855) until sequencing results were confirmed. A full list of primers used can be found in Supplementary Table 1.

### Human pluripotent stem cell culture and cardiac differentiation

Standard maintenance and passaging of hiPSCs was done as published previously.^34,35^ hiPSCs were differentiated into hiPSC-derived cardiomyocytes (hiPSC-CMs) using a standard monolayer Wnt-on and Wnt-off protocol.^36^ hiPSCs were cultured on Matrigel-coated (1 mg Matrigel aliquot in 24 mL of DMEM/F-12 medium, Corning, 356234) plates in mTeSR Plus medium (STEMCELL Technologies, 100-0276) and maintained for 3 days or until achieving 70–90% confluency. Differentiation was initiated by switching to RPMI 1640 basal medium (Gibco, 11875093) supplemented with 2% B27 without insulin (RPMI B27-; Gibco, A1895601) and 12 µM CHIR99021 (STEMCELL Technologies, 72054) for 24 h to activate Wnt signaling via GSK3 inhibition (Days 0-1). On day 3, cells were treated with 5 µM IWP4 (Tocris, 5214), a Wnt inhibitor, in RPMI B27- for 48 hours. From day 7 onward, the medium was switched to RPMI 1640 supplemented with 2% B27 with insulin (RPMI B27+; Gibco, 17504044), with media changes every 2-3 days. Spontaneous beating of hiPSC-CMs was typically observed between days 8 and 11. hiPSC-CM purification was performed using sodium L-lactate (Sigma Aldrich, 71718) in glucose-free RPMI B27+ (Gibco, 11879020) basal medium for 4 days. Purified hiPSC-CMs were then replated and subjected to a 3-week fatty-acid rich metabolic maturation protocol based on Feyen *et al.*,^37^ after which hiPSC-CMs were used for downstream assays. All downstream assays were performed with a minimum of 3 independent differentiation batches.

### Flow cytometry

To verify pluripotency, control and E4146D^+/-^ hiPSC lines were assessed for the pluripotency extracellular markers SSEA-4 (BD Pharmingen, 560796) and TRA-1-60 (BD Pharmingen, 560193). hiPSCs were dissociated using ReLeSR (STEMCELL Technologies, 100-0484) and collected in mTeSR Plus (STEMCELL Technologies, 100-0276). hiPSCs were centrifuged and transferred to a V-bottom 96-well plate for staining. Viability was assessed using a live/dead marker (fixable viability dye eFluor-780, Invitrogen, 65-0865-14) prior to antibody staining. hiPSCs were incubated with the antibody mixture for 30 minutes at room temperature in the dark, with fluorescence minus one (FMO) controls included for each marker. Following two washes in PBS, hiPSCs were resuspended in FACS buffer and analyzed on the BD FACSymphony A1 flow cytometer. Marker expression was quantified in comparison to FMO controls to confirm pluripotency (Supplementary Figure 1). A list of antibodies can be found in Supplementary Table 2.

### Confocal and high spatiotemporal optical mapping

For low density plating, hiPSC-CMs were replated for onto polymer coverslips (Ibidi, 10502-ibiTreat) coated with a 150 µL centrally placed droplet of Matrigel diluted 1:6 in DMEM/F-12 (0.0167% w/v; Gibco, 12634010). hiPSC-CMs were dissociated using an enzymatic protocol with Collagenase B (1 mM; STEMCELL Technologies, 07439) and Accumax (STEMCELL Technologies, 07921). hiPSC-CMs were first incubated in Collagenase B for 15-30 minutes. After Collagenase B treatment, the enzyme was removed, and hiPSC-CMs were incubated in Accumax as a secondary digestive step for 5-10 minutes and stopped with RPMI B27+. After centrifugation, hiPSC-CMs were resuspended in RPMI B27+ and ROCK inhibitor (Y-27632, Biogems, 1293823), counted, and 50,000 hiPSC-CMs were seeded onto Matrigel-coated coverslips. hiPSC-CMs were incubated at 37°C for 1 hour before adding the final culture volume, followed by a full media change the next day. hiPSC-CMs were allowed to adhere and rest for up to 14 days before imaging.

To form confluent hiPSC-CM monolayers, a high-density plating protocol was performed. hiPSC-CMs were dissociated as described above and 500,000-750,000 hiPSC-CMs were seeded onto polymer coverslips (Ibidi, 10502-ibiTreat) coated with a 150 µL centrally placed droplet of Matrigel diluted 1:6 in DMEM/F-12 (0.0167% w/v). hiPSC-CMs were allowed to adhere and rest for up to 14 days before imaging.

On the day of optical mapping, culture medium was aspirated and replaced with RPMI B27+ containing 5 µM Calbryte 630 (AAT Bioquest, 20721), as described previously^34,35^ and hiPSC-CMs were incubated for 1 h at 37 °C. For dual optical mapping of membrane voltage, we added 1x FluoVolt dye mixed with Powerlead (Invitrogen, F10488) according to the manufacturer’s protocol and incubated the hiPSC-CMs for an additional 15-20 minutes at 37°C. Coverslips were then washed two times with pre-warmed standard Tyrode’s solution, which is composed of (in mM): 117 NaCl, 5.7 KCl, 4.4 NaHCO_3_, 1.5 NaH_2_PO_4_-H_2_O, 1.7 MgCl_2_, 10 Na-HEPES (C_8_H_18_N_2_O_4_S), 5 glucose, 5 creatine, 5 Na-Pyruvic acid, and 1.8 CaCl_2_ (all from Sigma-Aldrich). To prevent motion artifacts, 5 µM Mavacamten (MYK-461, Selleckchem, S8861) was added to Tyrode’s solution. Coverslips were then transferred to a magnetic imaging chamber (Live Cell Instrument, Chamlide EC) equipped with two electrodes for field stimulation and kept in warm Tyrode’s solution immediately prior to imaging.

Confocal Ca^2+^ imaging of sparsely plated hiPSC-CMs was performed using the Leica TCS SP8 Confocal Microscope resonant scanning (8 kHz) at 63x (oil immersion), with a capture rate of 28 frames per second at 512 x 512. hiPSC-CMs were recorded under spontaneous beating and with electrical stimulation using an external stimulator. Confocal Ca^2+^ data was analyzed using a custom-made, in-house *R* code. Filtering included baseline drift correction (2^nd^ degree polynomial correction) and a moving average smoothing filter (maximum 7×7).

High spatiotemporal resolution optical mapping of Ca^2+^ and membrane voltage for hiPSC-CM monolayers was performed on the SciMedia dual CMOS camera, high-speed optical mapping system (SciMedia, MiCAM05-N256). The system was coupled with a SPECTRA light engine (Lumencore) for excitation and an ESTM-9 multifunctional stimulator (SciMedia) to synchronize electrical pacing, light source control, and camera timing during acquisition. Imaging was performed in an environmental chamber maintained at 37 °C and 5% CO₂. Signals from FluoVolt and Calbryte 630 reporters were spectrally separated and simultaneously captured (voltage: 107 fps, pixel dimensions: 1212 x 1080; Ca^2+^: 277 fps, pixel dimensions: 256 x 256) on dual cameras, with data acquisition and processing conducted using bV Workbench software (SciMedia) under both spontaneous and paced conditions. All analyses, including all ROI-based and tissue pixel-based mean and median quantifications, were conducted using the bV Workbench software (version 4.0) according to manufacturer guidelines. For all analyses, monolayers with abnormal or re-entrant spontaneous conduction patterns at baseline were excluded and not imaged further. Filtering included a 2D deconvolution mean filter, baseline drift correction (2^nd^ degree polynomial correction), and moving average filter (maximum 7×7, for Ca^2+^ data only). For quantitative analysis, only high-quality fluorescence signals (traces discernible with minimal background noise after filtering) were analyzed. One to two regions of interest per monolayer were analyzed for quantitative analyses, based on intra-monolayer variability. Tissue APD_90_, CaTD_80_, and activation maps were generated using the bV Workbench software. To test the efficacy of flecainide, hiPSC-CMs were incubated with Tyrode’s solution containing 1 µM flecainide (Sigma-Aldrich, F6777) at 37°C for 15 minutes before imaging.

### Programmed electrical stimulation protocols

For all programmed electrical stimulation protocols, we used the following pulse parameters: pulse width of 10 ms, amplitude of 5 mV, current of 60 mA, and biphasic. The long burst (LB) consisted of 25 consecutive, fixed interval stimuli (S1) at various frequencies/cycle lengths (e.g., 2 Hz, 500 ms basic cycle length). We tested LB protocols up to 5 Hz in some samples, based on how well each monolayer entrained. The long pause followed by a stimulus (LP) consisted of a 1000-1500 ms pause followed by an S1. The interval of the short-coupled stimulus (S2) following LP was started at an interval of 800 ms. Based on whether the S2 mimicked an EAD or not, we altered the interval until the S2 started approaching the tail end of the previous impulse. Then, the S2 interval was tested in decreasing 20 ms increments until there was complete loss of capture (which typically occurred around 200-280 ms). This was done to avoid pacing fatigue and phototoxicity given that each recording exposes hiPSC-CMs to 20-30 seconds of light. Typically, the signal would start to attenuate after completion of a full LBLPS protocol, which limited our ability to test the full LBLPS protocol at various frequencies in each monolayer.

### Multielectrode array

Field potential recordings were obtained using a Maestro Pro multielectrode array (MEA) system (Axion BioSystems, 62511) to assess beat period irregularity, field potential duration, and conduction velocity in isogenic control and E4146D^+/-^ hiPSC-CM monolayers. Prior to replating, 24-well or 48-well CytoView MEA plates (Axion BioSystems, M384-tMEA-24W and M768-tMEA-48B) were coated with Matrigel (1mg aliquot diluted in 6 mL DMEM/F-12 medium), adding a 12 µL droplet per well. Sterile ddH₂O was added to the outer rim of each plate to maintain humidity, and the plates were incubated at 37°C for 1-2 hours prior to replating. hiPSC-CMs were dissociated using the enzymatic protocol with Collagenase B and Accumax described above. hiPSC-CMs were seeded onto the MEA plate at a density of 150,000 cells/well. hiPSC-CMs were maintained in RPMI B27+, with ROCK inhibitor for the first day only, and the media was refreshed every 3 days thereafter. Replated hiPSC-CMs were allowed to adhere and rest for up to 14 days prior to data acquisition. Recordings were obtained at spontaneous beating, for ∼90 seconds per measurement, at 37°C and 5% CO₂. The AxIS Navigator Software (Axion BioSystems) was used for real-time monitoring of hiPSC-CM activity, and the Cardiac Analysis Tool (Axion BioSystems) was used for data analysis and processing. Field potential durations were corrected for beating rate using the Fridericia method.

### Immunocytochemistry

Immunocytochemistry was performed in hiPSC-CMs replated and cultured on micropatterned plastic coverslips (Curibio, ANFS-CS25) for up to 14 days to assess RyR2 expression and subcellular localization. After differentiation, metabolic selection, and three weeks of metabolic maturation, hiPSC-CMs were first washed with PBS and then fixed with 2% paraformaldehyde in PBS for 10 minutes at room temperature with gentle agitation. Following fixation, hiPSC-CMs were washed three times with PBS and incubated in 0.1 M glycine (Sigma-Aldrich, G2879) in PBS for 10 minutes, followed by permeabilization with 0.1% Triton X-100 (Sigma-Aldrich, X100) in PBS for 15 minutes and three subsequent washes with PBS. Non-specific binding was blocked using BlockAid blocking solution (Invitrogen, B10710) for 1 hour at room temperature. hiPSC-CMs were incubated overnight at 4 °C in a humidified chamber with a mouse monoclonal anti-RyR2 primary antibody (Invitrogen, MA3-916) diluted 1:150 in BlockAid solution. After three washes, samples were incubated with a goat anti-mouse Alexa Fluor™ 488-conjugated secondary antibody (Invitrogen, A11001) diluted 1:500 in BlockAid solution for 1-1.5 hours at room temperature in the dark. Nuclei were counterstained with Hoechst 33342 (Invitrogen, H3570) for 15 minutes at room temperature, followed by three final washes with PBS. Coverslips were mounted onto glass slides using ProLong Gold antifade mounting medium (Invitrogen, P10144) and sealed with transparent nail polish. Mounted samples were stored at 4 °C in the dark until imaging. Images were acquired using a Nikon Ti2 CSU-X1 spinning disk confocal microscope with a 60x water immersion objective and deconvolution was performed using Huygens Essentials software (Scientific Volume Imaging) default settings.

### RNA extraction and quantitative polymerase chain reaction

Control and E4146D^+/-^ hiPSC-CMs maintained in maturation media for 3 weeks were lysed, followed by RNA extraction and purification using the Qiagen RNA extraction and purification kit (Qiagen, 74104) according to the manufacturer’s protocol. cDNA was synthesized from 1000 ng of total RNA using a reverse transcription kit (Qiagen, 205311) according to the manufacturer’s protocol. Quantitative PCR was performed using a SsoAdvanced SYBR Green-based master mix (Bio-Rad, 1725271). Gene expression was quantified using the ΔΔCt method. Expression of *RYR2* was normalized to the cardiac-specific reference gene *TNNT2*. A list of primers used is shown in Supplementary Table 1.

### Mass-spectrometry proteomics

Control and E4146D^+/-^ hiPSC-CMs maintained in maturation media for 3 weeks were collected using a cell scraper and pelleted by centrifugation. Around 1,000,000 cells per replicate were lysed in 100 mM HEPES (pH 8.5) with 2% SDS and 1x Halt protease inhibitor cocktail (ThermoScientific, 78438). Lysates were heated at 95 °C for 5 min, cooled on ice for 3 min, and stored at-80 °C until proteomic analysis. Bottom-up mass spectrometry (MS) was performed as previously described.^38^ Briefly, lysates were sonicated and treated with benzonase (Novagen, 707463) to fragment chromatin, reduced with 10 mM Dithiothreitol, and alkylated with 50 mM chloroacetamide. Proteins were processed using the SP3 protocol, with hydrophilic and hydrophobic SpeedBeads magnetic carboxylate beads (50 mg/mL; Cytiva, GE45152105050250) and automation on a Thermo KingFisher Apex platform (Thermo Scientific) for high reproducibility. Proteins were bound to beads in 80% ethanol, washed with 90% ethanol, and digested on-bead with trypsin/Lys-C (enzyme/protein 1:75, Promega, V507A) in 100 mM ammonium bicarbonate for 16-18 h at 37 °C. Peptides were desalted and purified on Evotips prior to LC-MS analysis. Mass spectrometry was performed on the Ultra timsTOF (Bruker) coupled to an EvoSep liquid chromatography system, with peptides separated on a C18 analytical column (15 cm × 150 µm, 1.9 µm, Evosep, EV1106) using a 44-minute gradient. Full MS scans (100-1700 m/z) were collected alongside DIA acquisition windows. Raw data were analyzed in Spectronaut Pulsar X (Biognosys, version 19.2) using a human UniProt FASTA database and iRT peptide sequences. Search parameters included trypsin/P digestion, fixed carbamidomethylation (C), variable acetylation (protein N-term) and oxidation (M), peptide lengths 7-52 amino acids, and up to 2 missed cleavages. Protein and precursor FDR were set to 1%, and quantification required a minimum of 2 peptides per protein.

Resultant protein intensities were analyzed using Perseus (version 2.1.5.0). Intensities were log_2_-transformed, and differential protein expression between control and E4146D^+/-^hiPSC-CMs was conducted in Perseus using a 5% false discovery rate (FDR).

### Statistical analysis

Data points are presented as mean ± SD or median, as specified in figure legends. Statistical comparisons between two groups were performed using the Kruskal-Wallis test for continuous data, and the Chi-square test for categorical data. Pearson correlation was used for fold change correlation analyses. A two-sided *P* value < 0.05 was considered significant, except we applied a Benjamini–Hochberg false discovery rate (FDR) correction method to set the significance threshold for proteomics analyses. All statistical analyses were performed in GraphPad Prism, except for omics data for which we used Perseus.

## RESULTS

### Generation and molecular characterization of CRDS RyR2-E4146D^+/-^ hiPSC-CMs

We sought to investigate a CRDS-linked *RYR2* variant that was (1) feasible to genome edit into hiPSCs using CRISPR-Cas9, (2) characterized clinically with a presentation inconsistent with CPVT, (3) and functionally validated as a loss-of-function (LOF) variant *in vitro*. We identified the RyR2 variant p.E4146D as a promising candidate, which was first identified by Hirose et al. in a 2-month-old female infant.^5^ The patient was initially diagnosed with LQTS, however, continued to experience ventricular fibrillation (VF) and short-coupled variant of torsade de pointes (scTdP) despite optimal medical therapy for LQTS. This patient also had repolarization abnormalities suggestive of CRDS. This variant was confirmed to reduce RyR2 channel activity in HEK293 cells,^5^ and is structurally located within the U-motif of the central domain, which is critical for channel gating through its interaction with the C-terminal domain of the S6 helix.^39^

To investigate this variant, we generated E4146D^+/-^ (c.12438G>C, silenced PAM site via c.12441G>A) and isogenic control hiPSC lines (Figure 1A). Both hiPSC lines showed high expression of pluripotency markers SSEA-4 and TRA-1-60 (Supplementary Figure 1A-D). Cardiac differentiation of hiPSC lines was performed using a standard Wnt-signaling based hiPSC-CM differentiation protocol, followed by metabolic selection using lactate and a 3-week metabolic maturation protocol,^37^ after which hiPSC-CMs were used for downstream assays (Figure 1B).

**Figure 1.**
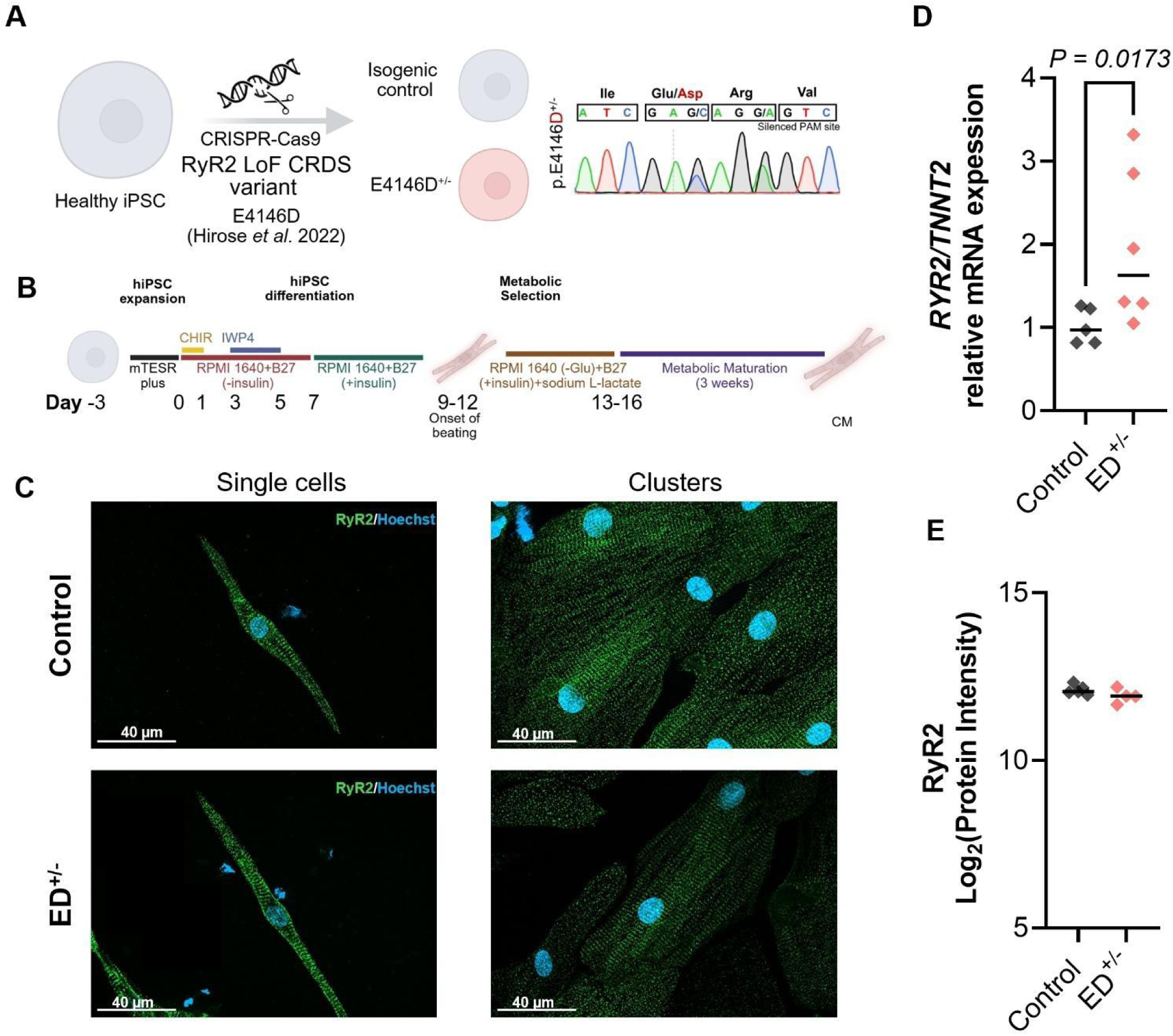
Generation and molecular characterization of RyR2-E4146D^+/-^ hiPSC-CMs. A) A commercially obtained hiPSC line (STAN248i-6717C1) from a healthy control was used to genome edit the CRDS-linked RyR2-E4146D variant using CRISPR-Cas9, generating a heterozygous E4146D^+/-^ hiPSC line. Sanger sequencing confirmed the edit and the silenced PAM site. B) Differentiation protocol used to generate control and E4146D^+/-^ hiPSC-CMs, which comprises a standard Wnt-on and Wnt-off protocol, followed by metabolic selection and 3 weeks of metabolic maturation treatment. C) RyR2 staining in control and E4146D^+/-^hiPSC-CMs plated at a low and high density on micropatterned coverslips. D) *RYR2* mRNA expression relative to *TNNT2* expression in control (n=5, 3 independent differentiation batches) and E4146D^+/-^ hiPSC-CMs (n=6, 4 independent differentiation batches). E) Log_2_-transformed protein intensity of RyR2 in control (n=5, 3 independent differentiation batches) and E4146D^+/-^ (n=4, 3 independent differentiation batches) hiPSC-CMs derived from bottom-up mass spectrometry.

To validate the success of our hiPSC-CM maturation protocol and examine RyR2 expression and cellular distribution, we performed imaging of control and E4146D^+/-^hiPSC-CMs plated on micropatterned coverslips at low and high densities. Micropatterns allow for longitudinal and regular alignment of hiPSC-CMs, removing the confounding effects of variable hiPSC-CM shapes and organization.^40^ We observed no gross structural abnormalities and z-disc alignment of RyR2 in control and E4146D^+/-^ hiPSC-CMs, indicating appropriate maturity for hiPSC-CMs (Figure 1C). Importantly, RyR2 expression and z-disc alignment were also observed in metabolically matured hiPSC-CMs plated on unpatterned surfaces (Supplementary Figure 2) suggesting alignment is not absolutely required for hiPSC-CM maturation.

Comparison of mRNA expression revealed an increase in *RYR2* mRNA expression in E4146D^+/-^ hiPSC-CMs (Figure 1D). To examine protein expression of RyR2 and other excitation-contraction coupling related proteins, we performed bottom-up mass spectrometry. Unlike mRNA expression, RyR2 protein expression was unchanged between control and E4146D^+/-^ hiPSC-CMs (Figure 1E), suggesting that *RYR2* mRNA expression increase may be a compensatory response to reduced RyR2 activity that is not reflected on the protein level. Protein expression levels of other key excitation-contraction coupling proteins including SERCA2a, Ca_V_1.2, Na_V_1.5, CASQ2, NCX1, and Cx43 were also unchanged (Supplementary Figure 3A). Additionally, the expression of critical regulators of RyR2 function, including all detected subunits of CAMKII, PKA, PKG, and PP1-3, did not significantly differ (Supplementary Figure 3B-C). Lastly, we found no difference in protein expression of the pan-cardiac markers TNNT2 and MYH7 (Supplementary Figure 3D) and maturation indices (TNNI3/TNNI1 and MYH7/MYH6) (Supplementary Figure 3E).

### Individual E4146D^+/-^ hiPSC-CMs show irregular spontaneous Ca^2+^ transients

Previous studies on LOF RyR2 hiPSC-CMs reported a greater propensity for irregular spontaneous Ca^2+^ transients (SCaTs) in individual hiPSC-CMs or small clusters, with and without the addition of isoproterenol,^30–32^ which comprised both EAD-like (i.e., systolic) and DAD-like (i.e., diastolic) Ca^2+^ transients. To understand how individual E4146D^+/-^ hiPSC-CMs behave, we plated hiPSC-CMs at a low density and measured Ca^2+^ release at spontaneous beating (Supplementary Figure 4A-B). Consistent with some previous studies in LOF RyR2 hiPSC-CM models,^29,30^ E4146D^+/-^ hiPSC-CMs displayed a significantly (P < 0.0001) greater propensity for irregular SCaTs compared to control hiPSC-CMs, the majority (77.1% of all abnormally beating cells) of which were late-systolic irregular SCaTs, although some cells displayed both late-systolic and diastolic irregular SCaTs (Supplementary Figure 4C-E), suggesting abnormal SR Ca^2+^ handling during both systole and diastole in individual CRDS hiPSC-CMs.

### RyR2-E4146D^+/-^ hiPSC-CM monolayers do not show arrhythmias at baseline

Next, we probed electrophysiological parameters on a tissue level in hiPSC-CM monolayers, including spontaneous arrhythmias, spontaneous beating rate, transients for action potential (AP) and Ca^2+^, and conduction velocity (CV) (Figure 2). We plated control and E4146D^+/-^hiPSC-CMs at high densities to induce monolayer formation, and performed high spatiotemporal optical mapping with fluorescent dye-based dual-imaging of membrane voltage (V_M_) and intracellular Ca^2+^. Measurements were performed 7-14 days post-replating depending on spontaneous beating recovery timeline in utilized hiPSC-CM monolayers (Figure 2A). No notable arrhythmias were observed at baseline spontaneous beating or 1 Hz steady state (Figure 2B), suggesting irregular SCaTs in individual hiPSC-CMs rarely translate to tissue-level arrhythmias.

**Figure 2.**
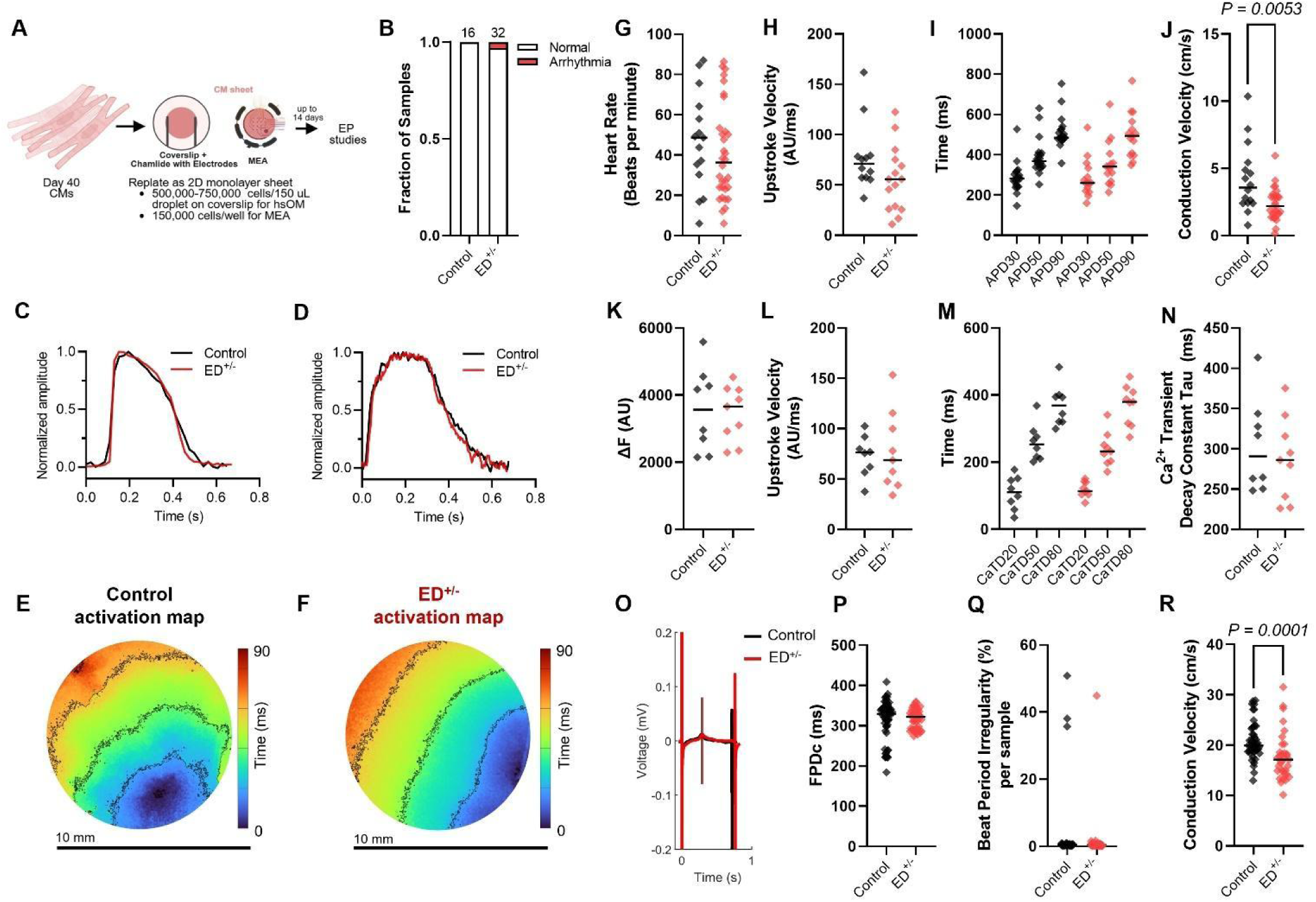
Baseline electrophysiological phenotyping of RyR2-E4146D^+/-^ and control cardiac monolayers. A) Workflow for replating day 40 hiPSC-CMs as monolayers for electrophysiological phenotyping using high spatiotemporal resolution optical mapping (hsOM) and multielectrode array (MEA). B) Fraction of control (n=16, 4 independent differentiation batches) and E4146D^+/-^ (ED^+/-^, n=32, 6 independent differentiation batches) monolayers displaying arrhythmias at baseline spontaneous or 1 Hz pacing. Only one arrhythmia was observed in 32 ED^+/-^ monolayers, which comprised one irregular, short-coupled beat after a normal spontaneous beat. C-D) Sample traces of C) membrane voltage measured using Fluovolt and D) Ca^2+^ transients measured using Calbryte 630 at 1.5 Hz steady pacing. E-F) Sample activation maps of E) control and F) ED^+/-^ monolayers. G) Spontaneous heart rate of control (n=15, 4 independent differentiation batches) and ED^+/-^(n=32, 6 independent differentiation batches) monolayers. H-I) Action potential H) upstroke velocity (n_control_=12, n_ED_=15, 4-5 independent differentiation batches) and I) duration at 30%, 50%, and 90% of repolarization (n_control_=19, n_ED_=15, 4-5 independent differentiation batches) at 1 Hz steady pacing. J) Tissue mean conduction velocity in control (n=16, 4 independent differentiation batches) and ED^+/-^ (n=29, 6 independent differentiation batches) monolayers. K-N) Ca^2+^ transient parameters including K) amplitude, L) upstroke velocity, M) transient decay at 20%, 50%, and 80%, and N) decay constant (Tau) in control (n=8, 4 independent differentiation batches) and ED^+/-^ (n=9, 5 independent differentiation batches) monolayers. For quantitative analyses of AP and Ca^2+^ transient parameters, 1-2 regions of interest (ROIs) were measured depending on intrasample tissue variability. O-P) MEA-based measurement of field potential duration, corrected for heart rate using the Fridericia method, in control (n=58, 3 independent differentiation batches) and ED^+/-^ (n=45, 3 independent differentiation batches) monolayers. Q) MEA-based measurement of the percentage of irregular beats in each sample in control (n=42, 3 independent differentiation batches) and ED^+/-^ (n=47, 3 independent differentiation batches) monolayers. R) MEA-based conduction velocity measurement in control (n=45, 3 independent differentiation batches) and ED^+/-^ (n=33, 3 independent differentiation batches) monolayers. Statistical comparisons between control and ED^+/-^ monolayers were performed using the Kruskal-Wallis test for all analyses, except a Chi-squared test in panel B.

Sample V_M_ and Ca^2+^ traces (Figure 2C-D) and activation maps (Figure 2E-F) for control and E4146D^+/-^ are shown. Spontaneous beating rate did not significantly differ between control and E4146D^+/-^ (Figure 2G). At 1 Hz steady-state pacing, we found no significant differences in AP parameters including upstroke velocity (Figure 2H) and AP durations at 30% (APD_30_), 50% (APD_50_), and 90% (APD_90_) of repolarization (Figure 2I). Interestingly, CV was significantly reduced in E4146D^+/-^ monolayers compared to control (Figure 2F).

Similar to AP parameters, there were no significant differences in Ca^2+^ transient parameters between control and E4146D^+/-^ monolayers, including Ca^2+^ transient amplitude (Figure 2K), upstroke velocity (Figure 2L), duration at 20% (CaTD_20_), 50% (CaTD_50_), and 80% (CaTD_80_) of decay (Figure 2M), and tau decay constant (Figure 2N).

Multi-electrode array (MEA)-based phenotyping of control and E4146D^+/-^ monolayers validated their similarity in heart-rate corrected field potential duration (FPDc) (Figure 2O-P), and lack of significant beat period irregularities at baseline (Figure 2Q). Consistent with optical mapping, CV was significantly (P < 0.001) reduced in E4146D^+/-^ relative to control monolayers (Figure 2R), corroborating our findings and suggesting a potential link between LOF RyR2 and reduced tissue CV.

### RyR2-E4146D^+/-^ hiPSC-CM monolayers are highly susceptible to pacing-induced alternans and re-entrant arrhythmias

The programmed electrical stimulation protocol shown to induce ventricular arrhythmias in CRDS patients and mouse hearts is composed of three consecutive parts: (1) a long burst of rapid pacing (LB), (2) a long pause (LP), and (3) a short-coupled extra stimulus (S) at varying coupling intervals. To better understand how CRDS and control hiPSC-CM monolayers would respond to each component, we employed the LBLPS protocol during high spatiotemporal OM with dual-imaging of V_M_ and intracellular Ca^2+^ and analyzed each component of the LBLPS protocol individually.

First, we applied a LB composed of 25 fixed interval stimuli (S1) at varying frequencies (2-5 Hz depending on entrainment capability of each monolayer) (Figure 3A). We found 25 beats to be sufficient to allow for entrainment of hiPSC-CMs monolayers, focusing on the last half of the LB pacing train for alternans analyses, as the first beats at any pacing frequency can alternate as the hiPSC-CMs adapt to faster beating frequencies. In control monolayers, rapid LB pacing typically resulted in spatially concordant alternans followed by global loss of entertainment (Figure 3B and D, top panel). Meanwhile, E4146D^+/-^ monolayers were more susceptible to the transition from spatially concordant to discordant alternans as pacing frequency increased (Figure 3C and D, bottom panel). Consistently, there was a significantly lower threshold for APD alternans and Ca^2+^ amplitude alternans (median 2 Hz) in E4146D^+/-^monolayers compared to controls (median 2.5 Hz; Figure 3E). The development of discordant alternans in E4146D^+/-^ monolayers resulted in notable spatial dispersion of repolarization, leading to unidirectional conduction blocks and re-entrant arrhythmias exclusively in E4146D^+/-^ monolayers (53.6%) (Figure 3F), while control monolayers remained unaffected. Sample V_M_ traces from a E4146D^+/-^ monolayer demonstrate the development of discordance, conduction blocks, and re-entry (Figure 3G, Supplementary Video 1). Re-entrant beats were most prominent at pacing frequencies which caused discordance but enabled sufficient diastolic intervals for re-entrant beats to propagate across the tissue before the following stimulus. Interestingly, while a few control monolayers eventually developed spatially discordant alternans, we did not observe any conduction blocks or re-entries in controls, suggesting a unique susceptibility to re-entrant arrhythmias in E4146D^+/-^ monolayers.

**Figure 3.**
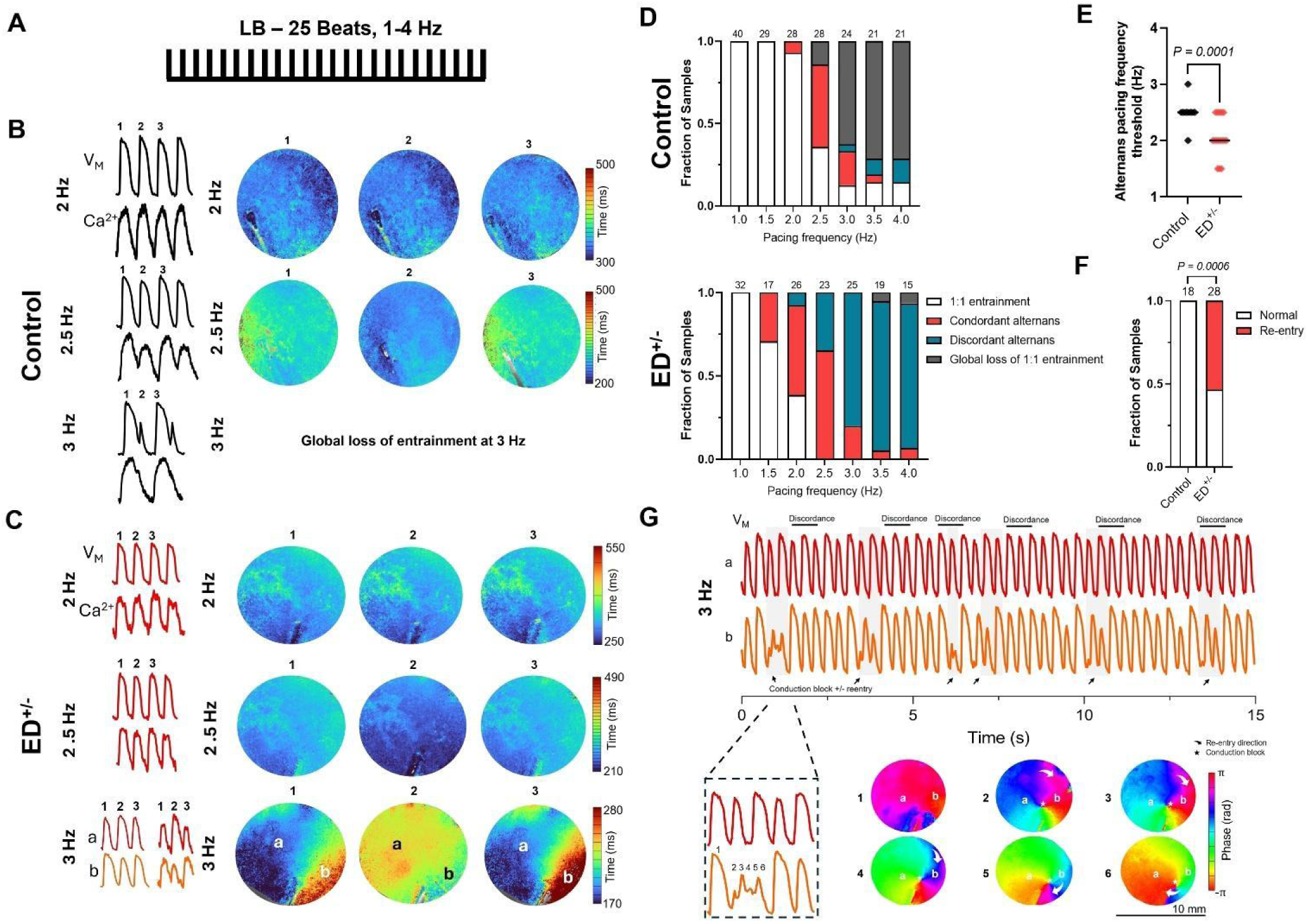
RyR2-E4146D^+/-^ cardiac monolayers are susceptible to pacing-induced alternans, conduction blocks, and re-entrant activation patterns. A) Long burst (LB) protocol applied as 25 consecutive, fixed-interval stimuli at various frequencies. B) Tissue action potential and Ca^2+^ transients and APD_90_ maps at 2 Hz, 2.5 Hz, and 3 Hz pacing for consecutive beats in control monolayers showing the development of concordant alternans at 2.5 Hz which do not transition to spatial discordance but instead experience a global loss of 1:1 capture at 3 Hz. C) Tissue action potential and Ca^2+^ transients and APD_90_ maps at 2 Hz, 2.5 Hz, and 3 Hz pacing for consecutive beats in E4146D^+/-^ (ED^+/-^) monolayers showing the tissue-level transition from spatially concordant to discordant APD and Ca^2+^ transient amplitude alternans. At 3 Hz, we show action potential and Ca^2+^ transient traces from two regions (a and b) which show spatially discordant alternans. D) Quantification of the fraction of control and ED^+/-^samples showing 1:1 capture, concordant alternans, discordant alternans, and global loss of 1:1 capture as LB pacing frequency is increased. The last half of the LB portion was used to determine alternans’ occurrence as the first few beats can alternate as cardiac tissue adapts to rapid pacing. E) Frequency threshold at which alternans are first observed in control (n=18, 4 independent differentiation batches) and ED^+/-^ (n=22, 5 independent differentiation batches) monolayers. F) Fraction of samples showing pacing-induced conduction blocks and re-entrant conduction patterns. G) Sample membrane voltage traces of a sample which demonstrates spatially discordant alternans leading to conduction blocks and re-entrant conduction patterns. The phase map first shows a normal beat, followed by a beat which demonstrated a re-entrant pattern due to a conduction block (indicated with a ★) in region b. Statistical comparisons were performed using the Kruskal-Wallis test for quantitative data, and the Chi-square test for categorical data.

### RyR2-E4146D^+/-^ hiPSC-CM monolayers display an abnormal post-pacing repolarization response unique to CRDS

A unique repolarization response after tachycardia or ventricular pacing has been observed in several patients with CRDS-linked LOF *RYR2* variants,^15,41^ including the patient carrying the here investigated E4146D^+/-^ variant.^5^ This response is marked by an abnormally large T wave amplitude and a prolonged QT interval on the first sinus beat post-tachycardia. This finding was first observed in members of a family possessing the RyR2-p.M4109R variant that had been diagnosed with CPVT prior to recognition of CRDS as a clinical entity.^41^ Subsequent realization that M4109R is a LOF variant led to the familial diagnosis being revised to CRDS and has resulted in the repolarization response being evaluated as a potential diagnostic tool for CRDS. A preliminary report involving a small number of CRDS patients and control groups revealed promising findings and concomitant studies involving a CRDS mouse model demonstrated a post-pacing increase in APD_90_ and Ca^2+^ transient amplitude that was unique compared to control and CPVT mouse models.^15^

To explore whether E4146D^+/-^ monolayers demonstrate this hallmark CRDS phenotype and better understand the impact of rapid pacing on the first spontaneous beat in human cardiac tissue, we used high spatiotemporal OM as described above and evaluated the first spontaneous beat post-LB compared to the rest. The same LB parameters (25 S1 stimuli, 2-5 Hz depending on entrainment capability of each monolayer) were applied (Figure 4A). An LP (800-1500 ms interval) was optionally applied if the monolayer did not return to spontaneous beating within 5 seconds. Sample traces demonstrate the tissue-level response to LB pacing in control (Figure 4B) and E4146D^+/-^ (Figure 4C) monolayers, with APD_90_ and CaTD_80_ maps shown for pre-pacing spontaneous, 1^st^ post-LB spontaneous, and 2^nd^ post-LB spontaneous beats. Interestingly, burst pacing induced an abnormal repolarization response in 35.7% of E4146D^+/-^ monolayers (Figure 4D). In all E4146D^+/-^ monolayers presenting with this phenotype, the first spontaneous beat was characterized by a significantly prolonged APD_90_ (mean 25.9% increase relative to pre-pacing), larger Ca^2+^ transient amplitude (mean 26.7% increase relative to pre-pacing) and CaTD_80_ (mean 41.7% increase relative to pre-pacing) (Figure 4E-G), with a positive and linear relationship (*R*^2^*=0.68, P* < 0.05) between APD_90_ prolongation and Ca^2+^ transient amplitude (Figure 4H). Interestingly, the abnormal beat was also characterized by a marked increase in the spatial dispersion of APD_90_ (*P* < 0.0001) and, frequently CaTD_80_ dispersion (*P* = 0.053), across each monolayer (Figure 4 I-J). This suggests an increase in repolarization heterogeneity in the first sinus beat post-pacing which is captured in our model, demonstrating the strong potential of this model in providing novel insights into the electrical vulnerability underlying the first sinus beat in CRDS.

**Figure 4.**
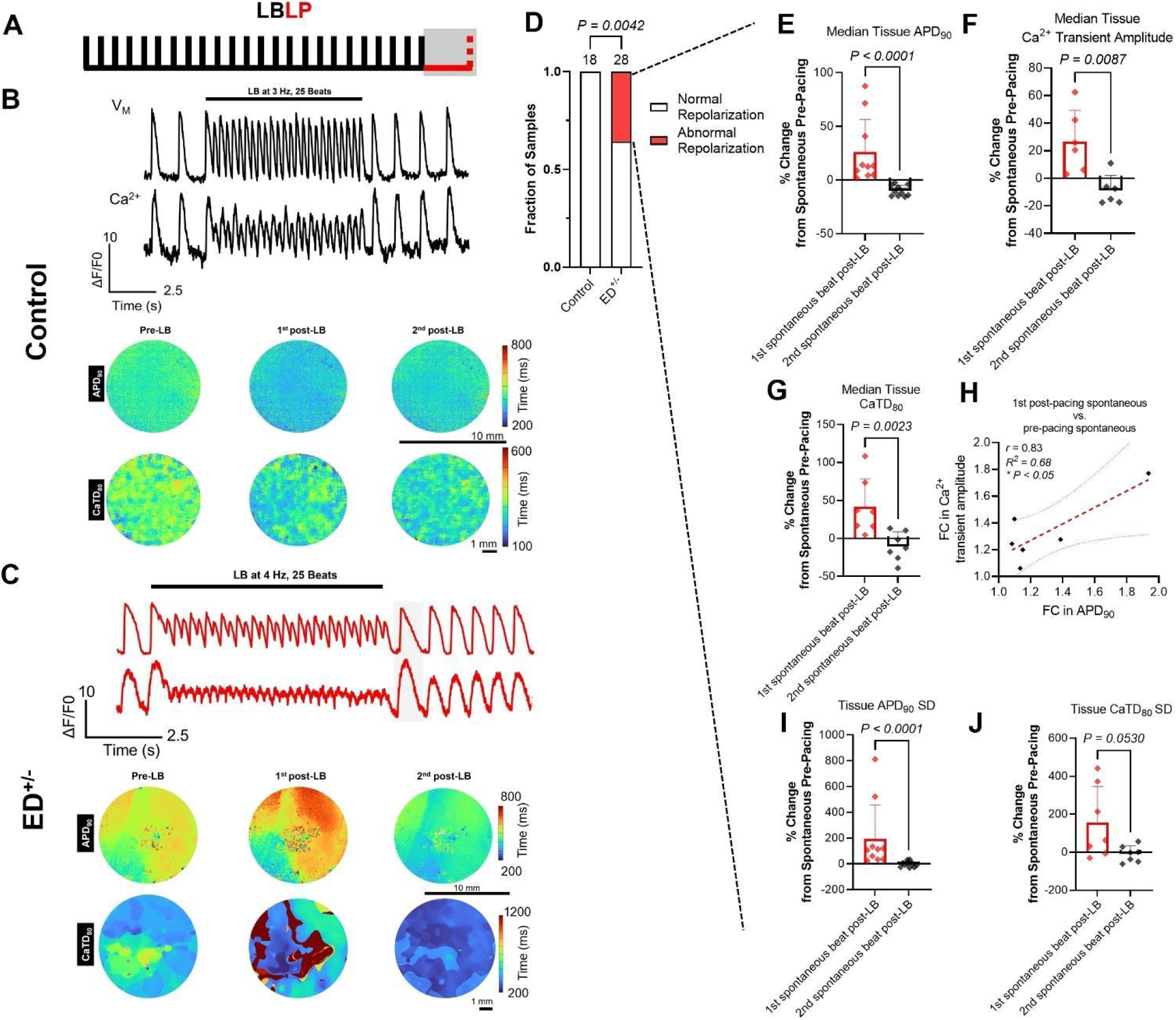
Post-pacing abnormal repolarization in RyR2-E4146D^+/-^ cardiac monolayers. A) A long burst (LB) consisting of 25 consecutive, fixed-interval stimuli at various frequencies was applied, followed by cessation of pacing to observe the post-pacing response of the first spontaneous beat. In rare cases, we applied an additional stimulus after 1000-1500 ms (LP) if the return to spontaneous beating exceeded 5 seconds. B) Representative membrane voltage and Ca^2+^ traces, tissue APD_90_ maps, and tissue CaTD_80_ maps, in control monolayers before, during, and after LB pacing, showing a return to normal spontaneous beating. C) Representative membrane voltage and Ca^2+^ traces, tissue APD_90_ maps, and tissue CaTD_80_ maps, in E4146D^+/-^ (ED^+/-^) monolayers before, during, and after LB pacing, showing an abnormal repolarization response in the first spontaneous beat post-pacing, characterized by a prolonged APD_90_ and a large Ca^2+^ transient amplitude. D) Fraction of control and ED^+/-^monolayers showing an abnormal repolarization response. E-G) Characterization of tissue-level changes in APD_90_ (n=10), Ca^2+^ transient amplitude (n=6), and CaTD_80_ (n=7), for the first abnormal spontaneous beat and the following spontaneous beat relative to average pre-pacing values. H) Correlation between the fold change in tissue APD_90_ post-pacing vs. pre-pacing and the fold change in Ca^2+^ transient amplitude post-pacing vs. pre-pacing for abnormal beats, showing a linear relationship between APD_90_ prolongation and larger Ca^2+^ transient amplitude. I-J) Quantification of the standard deviation (SD) of APD_90_ and CaTD_80_ values across each monolayer, representing the dispersion of APD_90_ (n=10), and CaTD_80_ (n=7) values for the first abnormal spontaneous beat and the following spontaneous beat compared to average pre-pacing values. Statistical comparisons were performed using the Kruskal-Wallis test for quantitative data, the Chi-square test for categorical data, and Pearsons’ correlation for correlation analysis.

We also tested the LBLP protocol in individual hiPSC-CMs. Using an LB protocol alone, we found that single hiPSC-CMs often did not return to spontaneous beating until >5 seconds post-beating. Thus, we applied an LBLP protocol, with an LB ranging from 2-3 Hz for 25 beats followed by an LP after 1000-1500 ms (Supplementary Figure 5A). Consistently, we found that a LBLP protocol resulted in a markedly larger Ca^2+^ transient amplitude in E4146D^+/-^hiPSC-CMs compared to control CMs, suggesting a larger SR Ca^2+^ load after the LB which is released with the following beat (Supplementary Figure 5B-D). Notably, not all E4146D^+/-^hiPSC-CMs demonstrated the same response, with greater variability (coefficient of variation = 24.4%) compared to controls (coefficient of variation = 17.8%) (Supplementary Figure 5D), which may explain the lower % inducibility of abnormal repolarization observed, and possibly the increase in repolarization dispersion when the phenotype is detected.

### NSVT-like re-entrant beats are triggered by LBLPS in RyR2-E4146D^+/-^ hiPSC-CM monolayers

In CRDS patients and mouse models, the LBLPS protocol can induce a range of ventricular arrhythmias ranging from a few ectopic or non-sustained ventricular tachycardia (NSVT) beats to sustained ventricular fibrillation and sudden cardiac death.^1,2,33,42^ Specifically, the arrhythmias are observed right after the short-coupled stimulus.

We reviewed traces from the implanted cardioverter-defibrillator in the E4146D^+/-^patient which captured the onset of ventricular fibrillation at 4 months old (Figure 5A). The tracing demonstrated a trigger which resembled many aspects of the LBLPS protocol. Specifically, during sinus tachycardia, a premature ventricular contraction (PVC) caused a compensatory pause and the following sinus beat demonstrated abnormal repolarization, marked by an abnormally tall T wave and prolonged QT interval. Then, the second sinus beat was followed by another PVC and subsequent polymorphic ventricular tachycardia/fibrillation.

**Figure 5.**
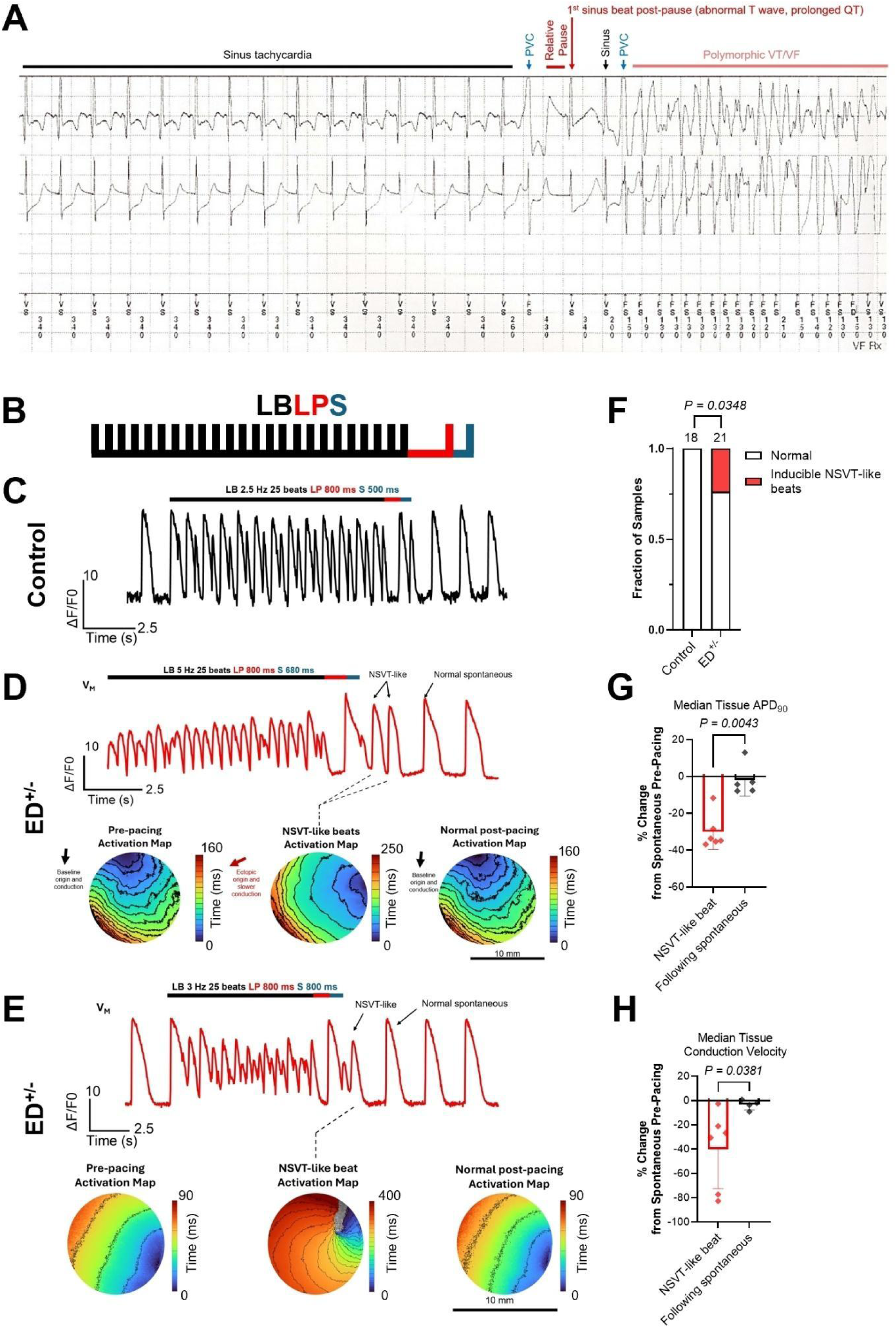
LBLPS induces pro-re-entrant, NSVT-like triggered activity in RyR2-E4146D^+/-^cardiac monolayers. A) Capture of ventricular fibrillation onset in the E4146D^+/-^ (ED^+/-^) patient at 4 months old using an implantable cardioverter-defibrillator. Trace shows a run of sinus tachycardia followed by a PVC which caused a compensatory pause. The following sinus beat displayed an abnormal T wave with a prolonged QT interval, followed a sinus beat and a subsequent PVC which likely triggered polymorphic ventricular tachycardia/fibrillation. B) The LBLPS protocol consists of a long burst (LB) composed of 25 consecutive, fixed interval stimuli applied at various frequencies, followed by a long pause (800-1500 ms) ending with a stimulus (LP), followed by a short-coupled extra stimulus (S) ranging from 800 ms to 200 ms to simulate an EAD. C) Sample membrane voltage trace showing a return to normal spontaneous beating after LBLPS with no triggered activity induced by LBLPS in control monolayers. D-E) Sample membrane voltage traces and activation maps in two ED^+/-^monolayers, one (D) showing two short-coupled, spontaneously triggered, NSVT-like beats induced by LBLPS which occur right after the S portion of the LBLPS protocol (which represents an EAD), whereby they show ectopic and slower conduction patterns compared to pre-pacing and normal post-pacing beats. Meanwhile, E) shows one short-coupled, spontaneously triggered, NSVT-like beat induced by LBLPS which shows a re-entrant and slower conduction pattern that is also ectopic compared to pre-pacing and normal post-pacing spontaneous beats. F) Fraction of control and ED^+/-^ monolayers which were inducible with LBLPS. G-H) Characterization of tissue median G) APD_90_ and H) conduction velocity for the spontaneously triggered, NSVT-like beats and the following normal spontaneous beat relative to average spontaneous pre-pacing values (n=6 NSVT beats). Statistical comparisons were performed using the Kruskal-Wallis test for quantitative data and the Chi-square test for categorical data. PVC: premature ventricular contraction.

To test for arrhythmia inducibility in our model, we applied an LBLPS protocol composed of: LB 25 beats at various frequencies (2-5 Hz depending on tissue entrainment), LP 800-1500 ms, and S ranging from 800 ms to 200 ms to simulate an EAD (Figure 5B). In controls, LBLPS did not induce any abnormal beats and the tissue returned to their normal spontaneous beating within a few seconds after pacing (Figure 5C). Notably, spontaneous beats most commonly originated from one specific region in each monolayer which remained the dominant pacemaker for the remainder of the recordings (data not shown). LBLPS was able to induce short-coupled, NSVT-like spontaneous beats right after the S in E4146D^+/-^monolayers, which was typically only 1 beat although one sample displayed two consecutive NSVT-like spontaneous beats (Figure 5D-E, Supplementary Videos 2-3). Inducibility was 0% in controls and 23.8% in E4146D^+/-^ (*P* < 0.05) (Figure 5F). These abnormal, NSVT-like triggered beats were characterized by an ectopic origin (compared to the spontaneous pacemaking region pre-pacing), shorter median tissue APD_90_ and CV (Figure 5G-H), and an often re-entrant conduction pattern (Figure 5D).

Although inducibility of NSVT-like triggered beats with LBLPS was modest in our model, we noted similar NSVT-like spontaneous beats during the LB portion at frequencies which resulted in spatially discordant alternans and EAD-like stimuli in some regions which differentially entrained, resembling a naturally occurring LBLPS-like sequence during discordant alternans (Figure 6A-B, Supplementary Video 4). These beats were similarly characterized by a re-entrant and ectopic activation pattern, and a significantly decreased median tissue APD_90_ and CV which did not differ from those triggered by programmed LBLPS (Figure 6C-D). Collective inducibility of NSVT-like beats right after LBLPS or during discordant LB pacing was 57.1% (Figure 6E). Notably, NSVT-like beats, even those induced after programmed LBLPS, were induced only when the LB was at a frequency which resulted in discordant alternans (Figure 6F), suggesting that dispersion of repolarization is critical for EAD-induced ectopic beats to initiate and/or propagate in CRDS human cardiac tissue.

**Figure 6.**
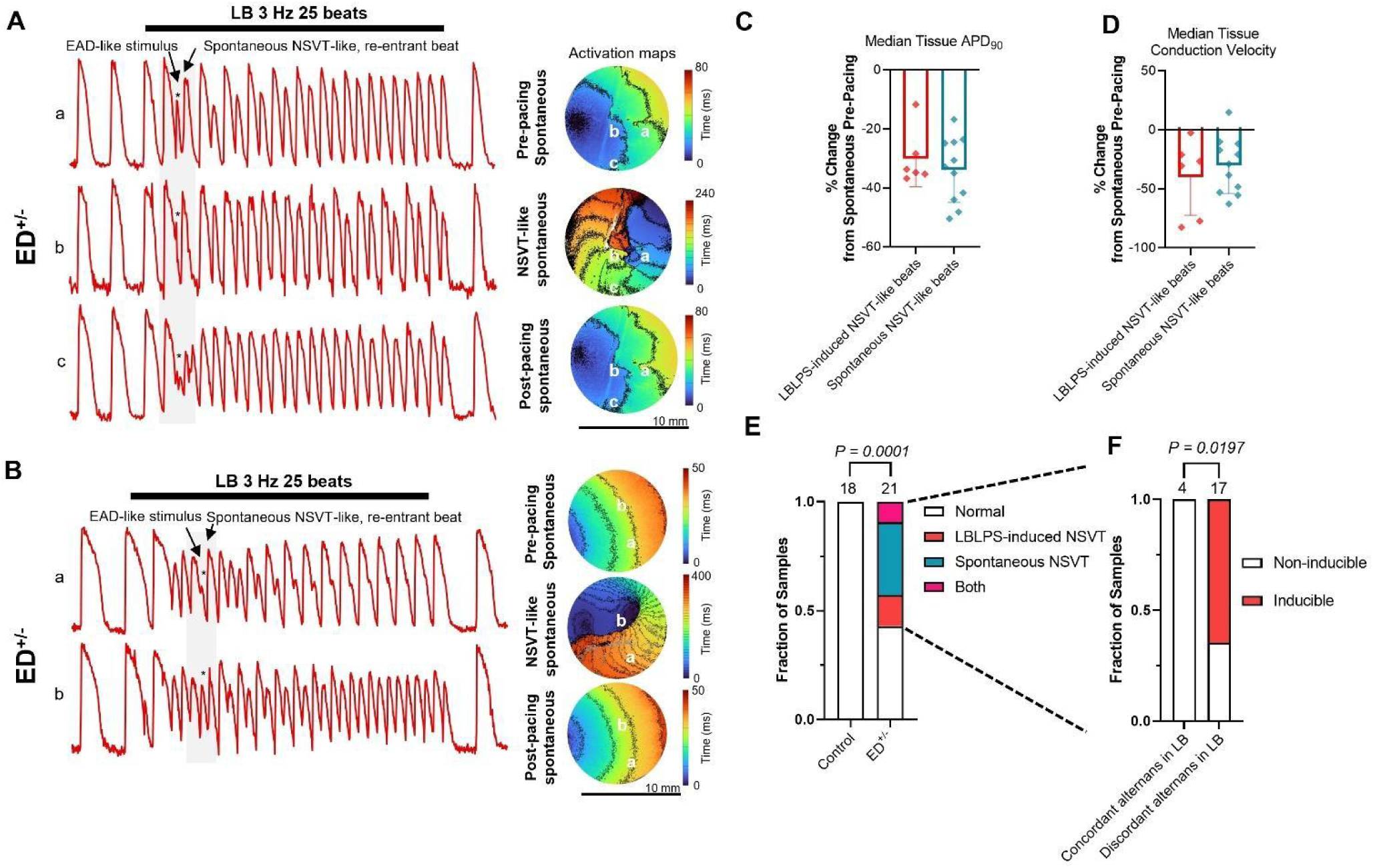
EAD-induced, NSVT-like re-entrant beats can develop spontaneously during spatially discordant alternans in RyR2-E4146D^+/-^ cardiac monolayers. A-B) Sample membrane voltage traces and activation maps in two E4146D^+/-^ (ED^+/-^) monolayers showing the development of spontaneously triggered, NSVT-like, re-entrant beats during pacing as a result of differential tissue entrainment, whereby the stimulus mimics an early after depolarization (EAD) in some regions but not others, suggesting that EADs during electrically heterogenous or vulnerable periods can trigger arrhythmogenic, re-entrant beats in ED^+/-^monolayers. C-D) Comparison of the tissue median APD_90_ and conduction velocity for LBLPS-induced, NSVT-like beats with spontaneously developed, NSVT-like beats relative to their average spontaneous pre-pacing values, showing similar characteristics of these abnormal beats (n=6 LBLPS-induced, n=11 spontaneously induced). E) Collective inducibility of LBLPS-induced and spontaneously developed NSVT-like triggered beats in control and ED^+/-^ monolayers. F) Fraction of ED^+/-^ samples which developed NSVT-like triggered beats when the long burst resulted in spatially concordant vs. discordant alternans. Statistical comparisons were performed using the Kruskal-Wallis test for quantitative data and the Chi-square test for categorical data.

### Flecainide alleviates arrhythmia inducibility in individual RyR2-E4146D^+/-^ hiPSC-CMs and monolayers

Evidence regarding the best treatment options for CRDS remains limited. Flecainide has shown promising results in alleviating ventricular arrhythmia burden in mouse models and CRDS patients who have undergone LBLPS testing.^1,2,33^ Moreover, administration of flecainide was shown to prevent the repolarization abnormality observed after ventricular pacing in a carrier of the LOF RyR2 p.M4109R variant.^41^ Relevant to our study, the E4146D^+/-^patient continued to experience arrhythmias until she was switched to a treatment regimen including carvedilol, flecainide, and an implantable cardioverter-defibrillator.^5^ Thus, we tested whether flecainide could rescue the inducible arrhythmias we observed in E4146D^+/-^hiPSC-CMs.

First, we tested the addition of flecainide on spontaneous beating in single hiPSC-CM, which significantly (P < 0.0001) reduced the number of cells with irregular SCaTs (Figure 7A). On a tissue level, flecainide did not affect the alternans threshold in E4146D^+/-^ monolayers (median threshold 2 Hz without and with flecainide) but reduced the likelihood of transitioning from spatially concordant to discordant alternans and instead promoted global loss of 1:1 entrainment at higher frequencies (Figure 7B-C), similar to control monolayers. Notably, flecainide significantly (P < 0.05) suppressed the abnormal repolarization response seen in E4146D^+/-^ monolayers (Figure 7D), and significantly (P < 0.05) reduced the inducibility of both spontaneous and LBLPS-induced NSVT-like triggered beats (Figure 7E). Thus, flecainide effectively protects against arrhythmia inducibility on both single cell and tissue levels, corroborating the limited evidence supporting its clinical efficacy. This demonstrates the utility of our model for future drug development for CRDS and other complex arrhythmias.

**Figure 7.**
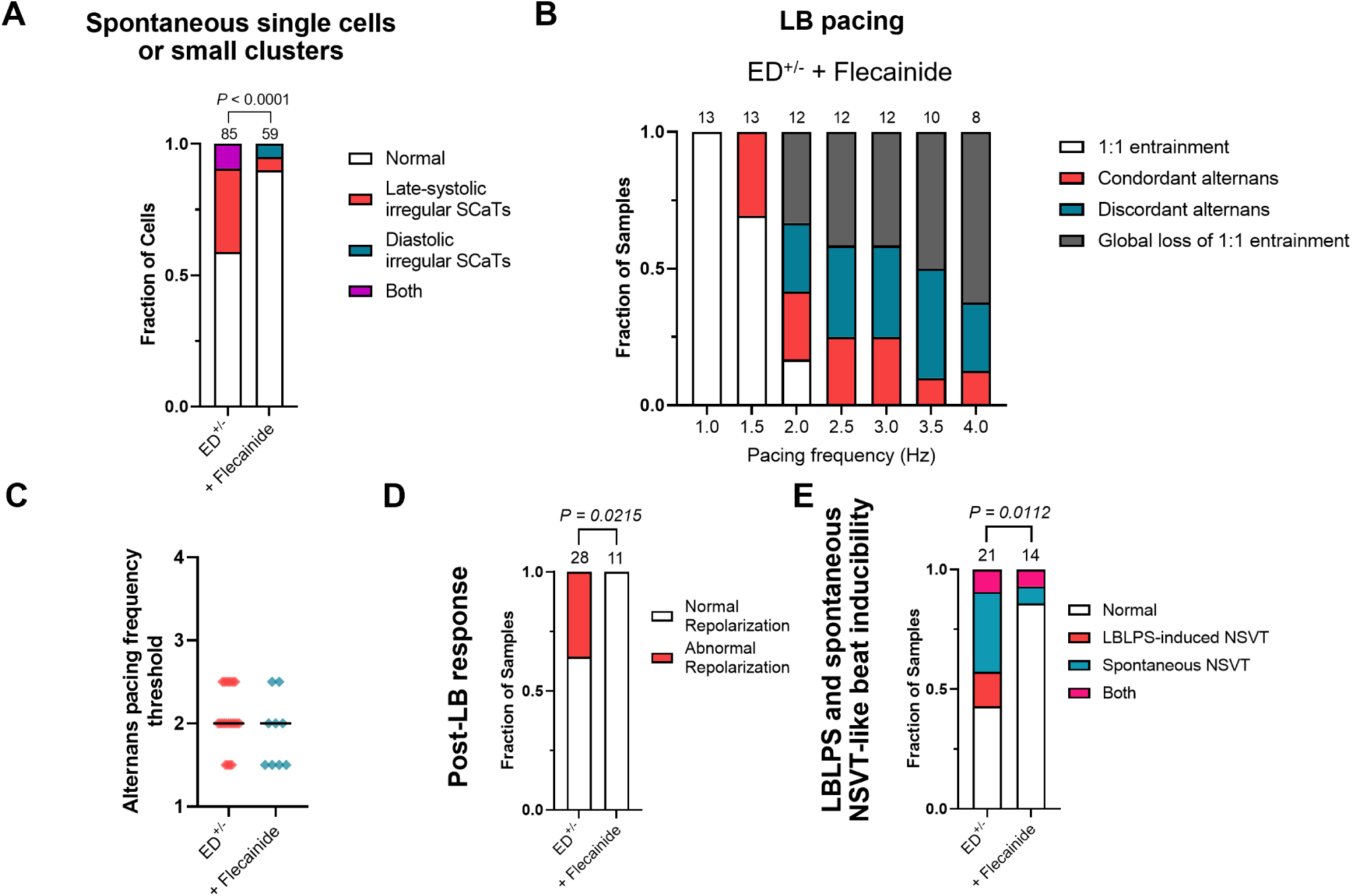
Flecainide alleviates arrhythmia burden and inducibility in RyR2-E4146D^+/-^ single hiPSC-CMs and cardiac monolayers. A) Fraction of single control and E4146D^+/-^ (ED^+/-^) hiPSC-CMs showing irregular spontaneous Ca^2+^ transients (SCaTs) without and with pre-treatment with flecainide (from 5-6 independent differentiation batches). B) Quantification of the fraction of d ED^+/-^ samples pre-treated with flecainide showing 1:1 capture, concordant alternans, discordant alternans, and global loss of 1:1 capture at LB pacing frequency is increased. The last half of the LB portion was used to determine alternans’ occurrence as the first few beats can alternate as cardiac tissue adapts to rapid pacing. C) Frequency threshold at which alternans are first observed in ED^+/-^ monolayers without (n=22, 5 independent differentiation batches) and with (n=9, 3 independent differentiation batches) pre-treatment with flecainide. D) Fraction of ED^+/-^ monolayers showing an abnormal repolarization response without and with pre-treatment with flecainide. E) Fraction of ED^+/-^ monolayers showing inducibility of LBLPS-induced and spontaneously induced NSVT-like triggered activity without and with pre-treatment with flecainide. Statistical comparisons were performed using the Kruskal-Wallis test for quantitative data and the Chi-square test for categorical data.

## DISCUSSION

Calcium release deficiency syndrome (CRDS) is a newly described inherited arrhythmia syndrome caused by LOF RyR2 genetic variants which remains much less understood compared to previously described channelopathies. Here, we present the first hiPSC-CM based CRDS model which recapitulates the key electrophysiological and clinical hallmarks of CRDS and provides key mechanistic insights into its pathogenic mechanisms. Using genome edited hiPSC-CMs carrying the RyR2-E4146D LOF variant which have been metabolically matured, we demonstrate that while arrhythmias are nearly absent on a tissue level at baseline, programmed electrical stimulation protocols can unmask clinically observed abnormalities such as post-pacing abnormal repolarization and short-coupled ectopy after LBLPS-like sequences in E4146D^+/-^ monolayers without beta-adrenergic stimulation. Moreover, our model provides compelling evidence of the EAD-mediated nature of triggered activity and re-entrant conduction during periods of electrical instability in CRDS, corroborating findings from mouse models,^1,14^ which have not been previously well recapitulated in previous hiPSC-CMs RyR2 LOF models.^30–32^

Normal RyR2 function and synchronised SR Ca^2+^ release are critical for rapid adaptation to increasing chronotropy. Previous work on RyR2 LOF mouse models supports the notion that reduced RyR2 expression or reduced function through LOF missense variants decrease the threshold for Ca^2+^ and APD alternans.^1,43–45^ Moreover, RyR2 LOF missense variants increase channel refractoriness,^1^ which is thought to be a critical driver of Ca^2+^ alternans.^46–48^ Consistently, we found that E4146D^+/-^ reduces the threshold for both Ca^2+^ and APD alternans in hiPSC-CM monolayers, promoting not only a lower threshold for alternans initiation but also the transition from spatially concordant to discordant alternans. The development of discordant alternans was critical for the generation of dispersed repolarization gradients, which are largely lacking at baseline in hiPSC-CM monolayers, creating vulnerable electrical periods during pacing which are susceptible to conduction blocks, re-entrant conduction, and EAD-mediated triggered beat initiation and propagation. While we did not measure RyR2 refractoriness, the susceptibility to discordant alternans in our model is likely driven by a global reduction in RyR2 refractoriness coupled with inter-cellular heterogeneity in RyR2 refractoriness thresholds due to the hetero-tetramer composition of each RyR2 complex (i.e., p.E4146D vs. wildtype RyR2) and inter-cellular differences in RyR2 expression. Clinically, discordant alternans can manifest as T wave alternans, which were reported in some traces in the E4146D^+/-^ patient.^5^ However, T wave alternans do not seem to be a consistent manifestation in many CRDS patients,^2^ suggesting clinical heterogeneity in the manifestation of RyR2 dysfunction. Interestingly, we also found a significant reduction in CV at spontaneous beating across E4146D^+/-^ monolayers, suggesting a potential role for SR Ca^2+^ release in mediating normal conduction. While the role of RyR2 in CV is not well understood, one group recently reported that gap-junction forming connexin 43 (Cx43) protein hemichannels are closely linked to and dependent on RyR2 for Ca^2+^-mediated activation.^49,50^ We found no difference in global Cx43 protein expression between control and E4146D^+/-^cardiomyocytes (Supplementary Figure 3A), suggesting the reduction is CV is likely not due to changes in total Cx43 protein expression. Importantly, the transition of alternans from spatially concordant to discordant could also be facilitated by slower CV in E4146D^+/-^monolayers, implicating both voltage and Ca^2+^ mediated mechanisms in electrical vulnerability.

The mechanism of arrhythmias caused by GOF RyR2 variants is generally well understood to be delayed-after depolarization (DAD)-mediated during periods of adrenergic stress.^10^ However, the mechanisms underlying LOF RyR2 variants remain much less clear. Some patients with CRDS present with arrhythmias related to exercise, but exercise stress testing does not reproduce arrhythmias like in CPVT. Meanwhile, other CRDS patients experience lethal arrhythmias at rest.^4^ Mouse models of RyR2 LOF have shown that isolated cardiomyocytes are susceptible to EADs and late-systolic, low amplitude Ca^2+^ oscillations, even without β-adrenergic stimulation.^1,14^ Moreover, the last portion of the LBLPS protocol, which has been shown to induce ventricular arrhythmias in several CRDS mouse models and patients,^1,33,42^ aims to simulate an EAD during the first post-pacing beat, suggesting that ventricular arrhythmias are triggered by EADs during electrically vulnerable states. The first hiPSC-CM study investigating LOF RyR2 missense variants utilized hiPSC-CMs with the M4109R variant, which at the time was thought to be a CPVT variant, found M4109R^+/-^hiPSC-CMs to be susceptible to some phase 3 EADs but mostly DADs, particularly with isoproterenol.^31^ This study, published in 2012, was a landmark study at the time, but the field of hiPSC-based disease modeling has rapidly advanced since then with large strides made in hiPSC-CM differentiation, maturation, and generation of isogenic controls using CRISPR-Cas9. A second study investigated a unique *RYR2* exon 1-4 homozygous duplication (RyR2-Dup) in a large Amish cohort with highly penetrant ventricular arrhythmias that did not fit typical CPVT criteria.^32^ The group generated hiPSC-CMs from two unrelated patients which showed significantly reduced RyR2 expression and caffeine-induced Ca^2+^ release compared to hiPSC-CMs from unrelated healthy individuals, suggesting a LOF phenotype. However, RyR2-Dup hiPSC-CMs were shown to be susceptible to DADs, particularly with 100 nM isoproterenol. This model may be a unique form of an RyR2 channelopathy, and it is unclear if its pathophysiology is reflective of the majority of reported CRDS cases. Another group generated two homozygous missense LOF RyR2 hiPSC-CM lines (Q3925E, E3848A), which demonstrated a notable propensity to late-systolic Ca^2+^ transients resembling the ones we observed here.^29^ Lastly, an hiPSC-CM study was conducted for the R4790Ter variant, which was recently shown to result in RyR2 LOF through a dominant-negative effect on WT RyR2.^51^ In another study this variant similarly demonstrated late-systolic Ca^2+^ transients which increased in propensity with 10 µM isoproterenol leading the authors to suggest that R4790Ter is a GOF variant.^30^

In our study, we leveraged metabolically matured hiPSC-CMs and focused on tissue-level mechanisms of RyR2 LOF and the clinically observed triggers, which we hypothesized would better recapitulate the arrhythmias CRDS patients experience. Our single hiPSC-CM Ca^2+^ imaging showed a higher propensity for late-systolic Ca^2+^ transients than diastolic Ca^2+^ transients at baseline, which may be representative of EADs. However, these did not translate to baseline arrhythmias on a tissue level. Although the E4146D^+/-^ patient was initially diagnosed with LQTS due to the presence of intermittently prolonged QT intervals after ventricular ectopy,^5^ we found no differences in heart rate, AP parameters, including APD_30-90_, or FPDc values between control and E4146D^+/-^ monolayers, at baseline or steady-state 1 Hz pacing. This is consistent with the lack of QT prolongation at baseline in the E4146D^+/-^ patient and other CRDS patients. We also found no differences in baseline Ca^2+^ transient parameters, including Ca^2+^ transient amplitude and decay time, suggesting potential compensatory mechanisms to maintain Ca^2+^ release during systole, such as enhanced *I*_Ca,L_, as previously shown in RyR2 LOF mouse and human CMs.^1,29^

Using carefully designed programmed electrical stimulation protocols which mimic the sequence of electrical events shown to trigger arrhythmias in intact CRDS mouse hearts and patients, we provide compelling evidence that EADs during electrically vulnerable periods are the trigger for arrhythmias in CRDS. We show specifically that the LBLPS sequence can trigger NSVT-like, ectopic triggered activity which favours re-entry in E4146D^+/-^ cardiac monolayers. Notably, however, the programmed LBLPS sequence was not always required for induced triggered activity as EADs during periods of increased dispersion of repolarization during the burst phase were also able to trigger spontaneous, re-entrant triggered beats. This is particularly important, as arrhythmias in the A4680G RyR2 LOF mouse model seem to develop without programmed LBLPS testing,^52^ but instead with a stress challenge including 200 nM isoproterenol and 3.6 mM Ca^2+^ Tyrode’s solution, indicating a link between increased SR Ca^2+^ load and arrhythmia vulnerability.^14^ Hence, several triggers may promote electrical vulnerability, during which EADs could lead to malignant, re-entrant arrhythmias. Our approach emphasizes the importance of maturation and tissue-level analyses in hiPSC-CM models for complex arrhythmias as seen in CRDS, as arrhythmias seen in single cells can be difficult to translate to the clinical phenotypes observed.

Importantly, we observed marked APD_90_ prolongation and a larger Ca^2+^ transient amplitude in the first beat after pacing in some E4146D^+/-^ monolayers and single hiPSC-CMs, explaining the clinical observation of abnormal repolarization post-pacing in several CRDS patients that is currently being tested as a clinical diagnostic test.^15^ This beat was characterized by spatially dispersed APD_90_, and to some extent CaTD_80_, prolongation across the tissue, rendering the first beat post-pacing particularly vulnerable to EAD-mediated arrhythmias. Given that the T wave amplitude is a marker of transmural and apico-basal repolarization heterogeneity in the intact heart,^53^ the pronounced enlargement of the T wave amplitude observed on the first sinus beat post-pacing in patients reflects an increased repolarization heterogeneity that is recapitulated in our *in vitro* system. While the reasons driving repolarization heterogeneity remain unclear, we found that E4146D^+/-^ hiPSC-CMs displayed a more variable Ca^2+^ release compared to control hiPSC-CMs after pacing, which could be due to the hetero-tetrameric RyR2 composition in heterozygous carriers and the subsequent variable Ca^2+^ release activity of each tetramer. Thus, our *in vitro* system successfully recapitulates the abnormal post-pacing repolarization response observed in CRDS patients, although inducibility remains relatively limited and the factors contributing to the inducibility of the phenotype in each monolayer remain unclear.

It remains unknown which therapeutics are most effective for CRDS patients, particularly given the low number of cases identified to date. Flecainide has shown the most promise thus far in preventing inducible arrhythmias with programmed electrical stimulation in M4109R^+/-^ and A4142T^+/-^ carriers,^33,41^ and the D4646A^+/-^ mouse model.^1^ Flecainide is known to be an effective agent for CPVT given alongside β-blockers, and is proposed to directly inhibit RyR2-mediated SR Ca^2+^ release in addition to its Na^+^ channel blocking effects.^54–56^ How flecainide would alleviate arrhythmia inducibility in CRDS remains unclear, however. Recently, a study generated a mouse model with combined silencing of all three canonical RyR2 phosphorylation sites (S2031A/S2808A/S2814A) that resulted in a CRDS-like phenotype including reduced Ca^2+^ leak and a greater propensity for phase 3 EADs, which also responded well to flecainide and lidocaine, implicating *I*_Na_ reactivation as a critical arrhythmogenic mechanism that could be suppressed by flecainide.^57^ We found flecainide to effectively reduce electrical vulnerability and arrhythmias on both single cell and tissue levels, consistent with the improvement observed in the E4146D^+/-^ patient, but whether this is driven by RyR2 inhibition, *I*_Na_ inhibition, or both, remains unknown.

## LIMITATIONS

There are several limitations in this study. First, while we mainly focused on tissue-level arrhythmia mechanisms, we did not investigate potential ion current remodeling or directly measured action potentials using patch clamp in single hiPSC-CMs. Moreover, the use of OM and fluorescent dyes, which can be cytotoxic over time, limited the ability to more extensively test programmed electrical stimulation protocols in each monolayer. Less invasive and higher throughput methods may be more suitable for high throughput drug screening in the future, now that we have a protocol to induce the phenotype. While we focused on one variant which is located in the U-motif of RyR2, which could limit the generalizability of our findings to LOF variants in other regions, the mechanisms seem largely consistent with C-terminal domain and transmembrane region variants which have been characterized thus far.^1,12,13,58^ Lastly, we focused on testing whether the efficacy of flecainide is recapitulated in our model, but further drug testing with other clinically available and experimental agents remains necessary.

## CONCLUSIONS

Collectively, we have developed the most robust hiPSC-CM model for CRDS to date, and the first to recapitulate key clinical hallmarks of the disease including programmed electrical stimulation-induced arrhythmias and repolarization abnormalities. Mechanistically, our model proposes EAD-mediated triggered activity during electrically vulnerable periods as the main mechanism of arrhythmia in CRDS, with LBLPS being one potential trigger given the marked repolarization heterogeneity of the first post-pacing beat. Notably, our study comprehensively details the electrical features of EAD-mediated triggered activity, providing novel mechanistic insights into how re-entry is initiated and propagated. Importantly, this human CRDS model has strong potential as a platform for targeted drug and gene therapy development, with flecainide being identified as an effective therapeutic in this study. Our approach may also provide novel mechanisms into other complex, tissue-level arrhythmias, such as short-coupled ventricular fibrillation, which remain largely elusive.^59^

### Author Contributions

S.F.D. and G.F.T. conceived the project. G.F.T. and M.P. supervised the project. S.F.D. designed and performed all experiments and analyses. A.A. and M.B. provided technical guidance and training. T.H. performed flow cytometry staining and analyses. S.O. contributed clinical data for the RyR2-E4146D patient. T.M.R., J.D.R., and S.S. provided clinical insights and ECG interpretation. F.J. and P.F.L. provided mass-spectrometry sample processing and protein quantitation services. F.V.P, R.A.R., L.H.M., E.D.W.M., and S.R.W.C. provided technical, experimental, and intellectual guidance. S.F.D. wrote the first manuscript draft. All authors contributed to manuscript revision. All authors approve the final version of the manuscript.

## Acknowledgements

The authors would like to thank the BC Children’s Hospital Research Institute (BCCHR) Imaging Core (Mr. Jingsong Wang) and Dr. Parisa Asghari for providing technical imaging support.

## Sources of Funding

This study was supported through funding from the Canadian Institutes of Health Research (CIHR) (GR020601 to G.F.T and F.V.P), the Mining for Miracles fund on behalf of the BC Children’s Hospital Foundation (to G.F.T and S.S.) and the Rare Diseases: Models and Mechanisms Network (to A.A. and M.P.). S.F.D. was financially supported through a CIHR Graduate Research Scholarship – Doctoral award (515168) and a Friedman MD/PhD studentship by the University of British Columbia. L.H.M. received funding from the Spanish Ministry of Science, Innovation and Universities (PID2023-152610OB-C21). R.A.R. received funding from CIHR (PJT203970 and PJT204002).

## Disclosures

T.M.R. reported receiving personal fees from Cardurion Pharmaceuticals. All other authors have no disclosures.

## ABBREVIATIONS

APD: Action potential duration
CPVT: Catecholaminergic polymorphic ventricular tachycardia
CRDS: Calcium release deficiency syndrome
DAD: Delayed afterdepolarization
EAD: Early afterdepolarization
GOF: Gain-of-function
hiPSC-CM: Human induced pluripotent stem cell-derived cardiomyocyte
IVF: Idiopathic ventricular fibrillation
LBLPS: Long burst, long pause, short-coupled extra stimulus
LOF: Loss-of-function
LQTS: Long QT syndrome
MEA: Multi-electrode array
NSVT: Non-sustained ventricular tachycardia
RyR2: Cardiac ryanodine receptor
scTdP: Short-coupled Torsades de Pointes
scVF: Short-coupled ventricular fibrillation
SOICR: Store overload-induced Ca2+ release
SR: Sarcoplasmic reticulum

