## Supplemental Materials for "Programmed electrical stimulation in human iPSC-derived cardiomyocytes reveals mechanisms of lethal arrhythmias in Calcium Release Deficiency Syndrome"

### Supplementary File Outline

Supplementary Table 1……….……………………………………………………………2

Supplementary Table 2……….……………………………………………………………3

Supplementary Figure 1……………………………………………………………………..4

Supplementary Figure 2……………………………………………………………………..5

Supplementary Figure 3……………………………………………………………………..6

Supplementary Figure 4……………………………………………………………………..7

Supplementary Figure 5……………………………………………………………………..8

**Supplementary Table 1.** List of primers and DNA sequences used in this study.

| Target | Forward (5'-3') | Reverse (5'-3') |
|---|---|---|
| ***Genome Editing*** | | |
| E4146D genomic region, amplification and sanger sequencing | GAGGGACTTCCACAAAGCGA | TTCTCTCCGCCTTCGTTGAC |
| sgRNA | 5'AAGCGCCAAACGCATCGAGA'3 | |
| ssODN | 5'TCTCCCACTGGGTTCGGCTGGACTCACTGATTTCAAAATAGACTCTGTCGATGCGTTTGGCGCTTCCCATGATTTCGATGCGGCCCAGAAAGGGCTGGA'3 | |
| ***Quantitative PCR*** | | |
| *TNNT2* | TTCACCAAAGATCTGCTCCTCGCT | TTATTACTGGTGTGGAGTGGGTGT |
| *RYR2* | CCTTGCCTGAGTGCAGTTG | TTGAGGTATCAACAGGTTGTGG |

**Supplementary Table 2.** List of antibodies used in this study.

| **Target** | **Antibody** | **Concentration** |
|---|---|---|
| ***Flow Cytometry*** | | |
| SSEA-4 | Mouse anti-SSEA-4 Alexa Fluor™ 647, BD Pharmingen, 560796, clone MC813-70, lot #3097806 | 1:50 |
| TRA-1-60 | Mouse anti-human TRA-1-60 PE**,** BD Pharmingen, 560193, clone TRA-1-60, lot #2048014 | 1:50 |
| Viability | eBioscience Fixable Viability Dye eFluor™ 780, Invitrogen, 65-0865-14 | 1:1000 |
| ***Immunocytochemistry*** | | |
| RyR2 | Primary: Mouse monoclonal anti-RyR2, Invitrogen, MA3-916, clone C3-33, lot# ZG397636 | 1:150 |
| | Secondary: Goat anti-mouse Alexa Fluor™ 488-conjugated secondary antibody, Invitrogen, A11001, lot# 2659299 | 1:500 |
| Nuclei | Hoechst 33342, Invitrogen, H3570 | 1:10,000 |

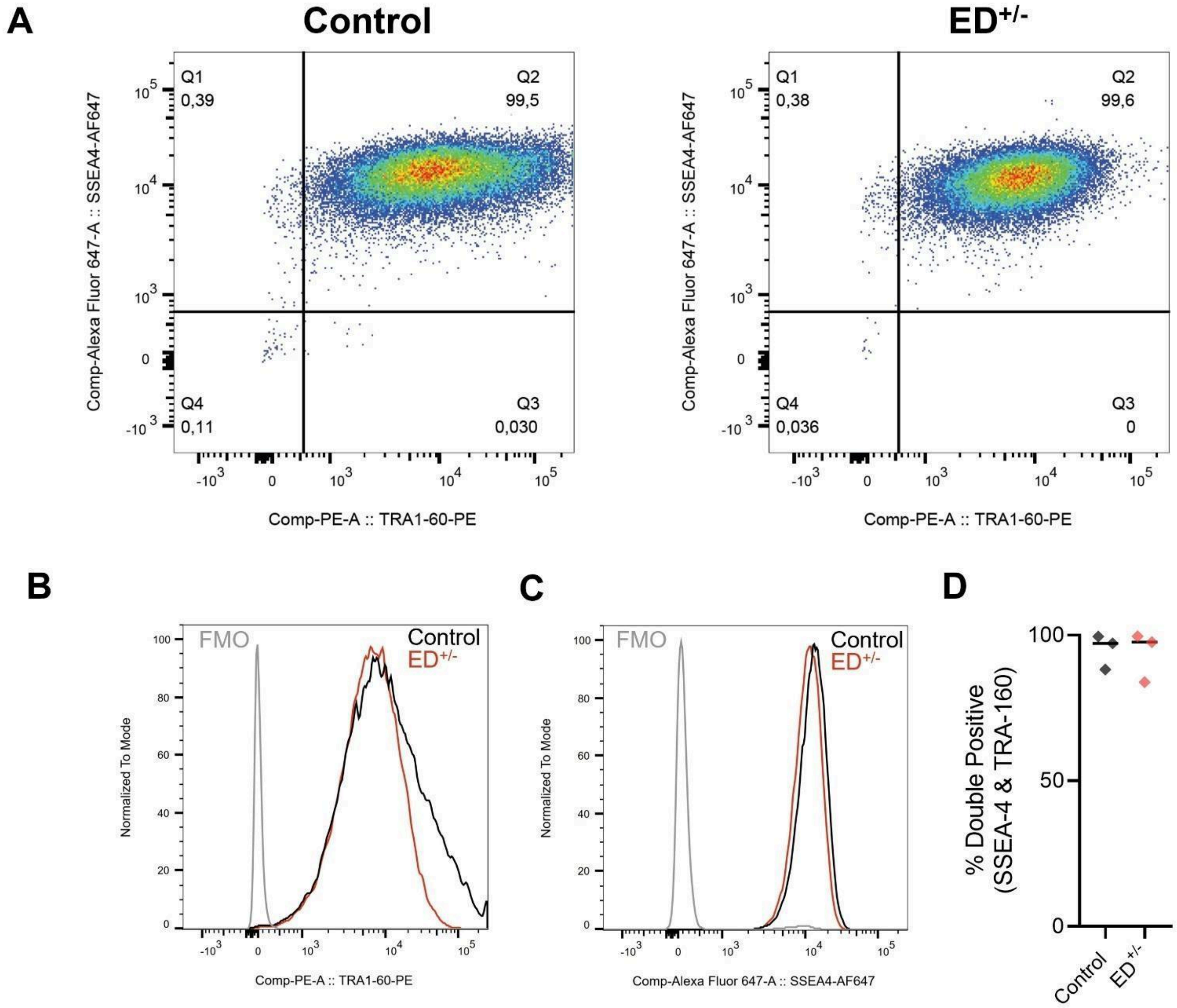


**Supplementary Figure 1.** Pluripotency assessment of control and E4146D$^{+/-}$ hiPSC lines. A) Flow density plot showing % of cells co-expressing pluripotency markers SSEA-4 and TRA-1-60. B-C) Sample histograms showing expression of B) TRA-1-60 and C) SSEA-4 in control and E4146D$^{+/-}$ (ED$^{+/-}$) hiPSCs compared to fluorescence minus one (FMO) controls. E) Percentage of hiPSCs co-expressing SSEA-4 and TRA-1-60. Each replicate represents an independent passage (n=3).

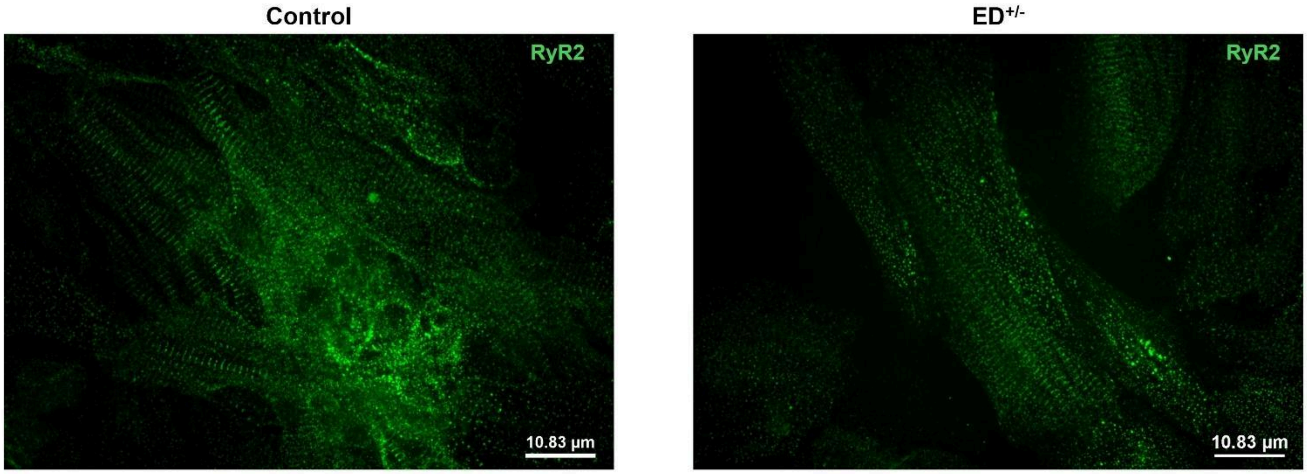


**Supplementary Figure 2.** Expression of RyR2 in hiPSC-CMs cultured on unpatterned coverslips. Sample RyR2 staining showing Z-disc alignment of RyR2 in control and E4146D$^{+/-}$ hiPSC-CMs cultured on unpatterned plastic coverslips after treatment with metabolic maturation media.

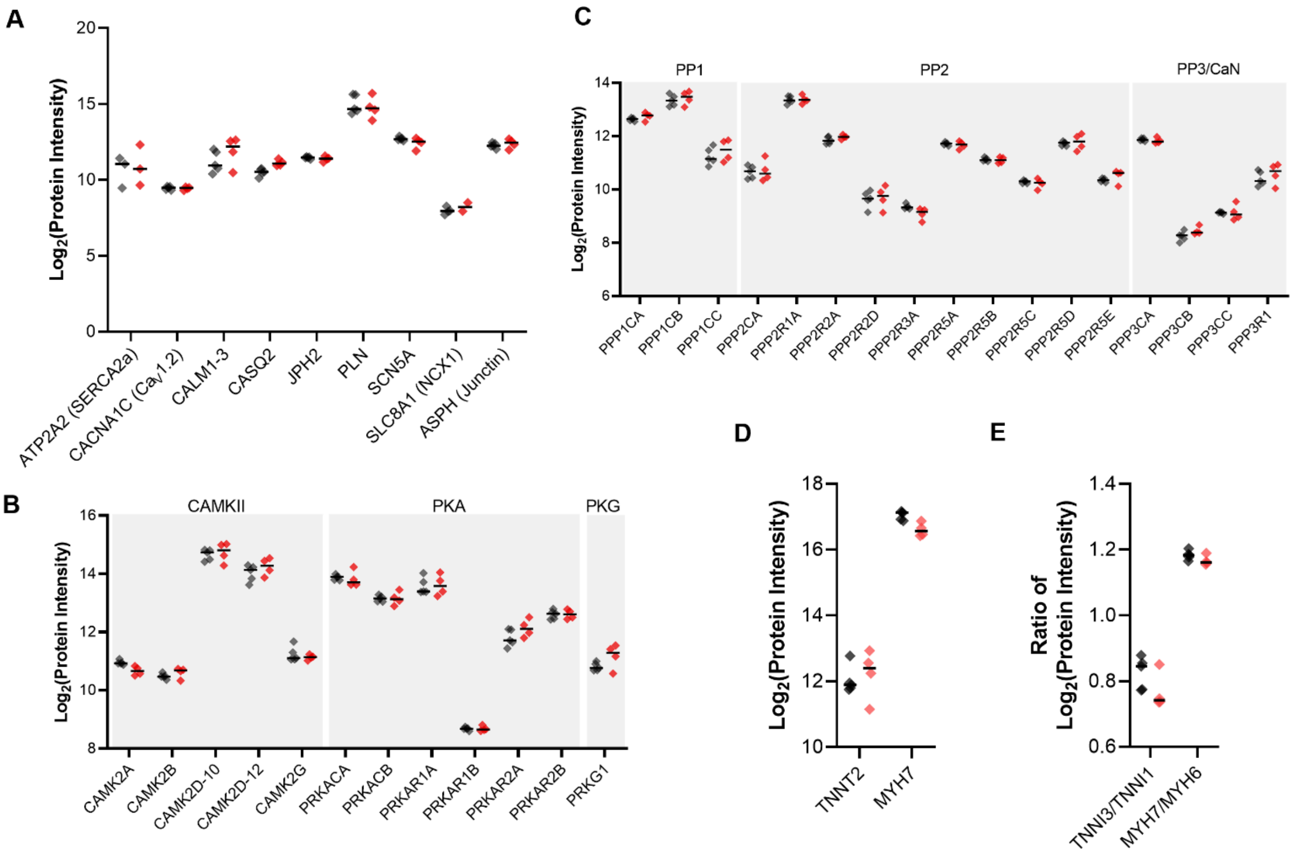


**Supplementary Figure 3.** Protein expression differences for excitation-contraction coupling proteins, RyR2 regulators, pan-cardiac markers, and maturation indicators. $Log_2$-transformed protein intensities of A) proteins involved in excitation-contraction coupling, B-C) kinases and phosphatases involved in the regulation of RyR2 and other relevant excitation-contraction coupling proteins, and D) pan-cardiac markers indicating similar cardiomyocyte composition in samples. E) Maturation indicators, including the ratio of TNNI3/TNNI1 and MYH7/MYH6.

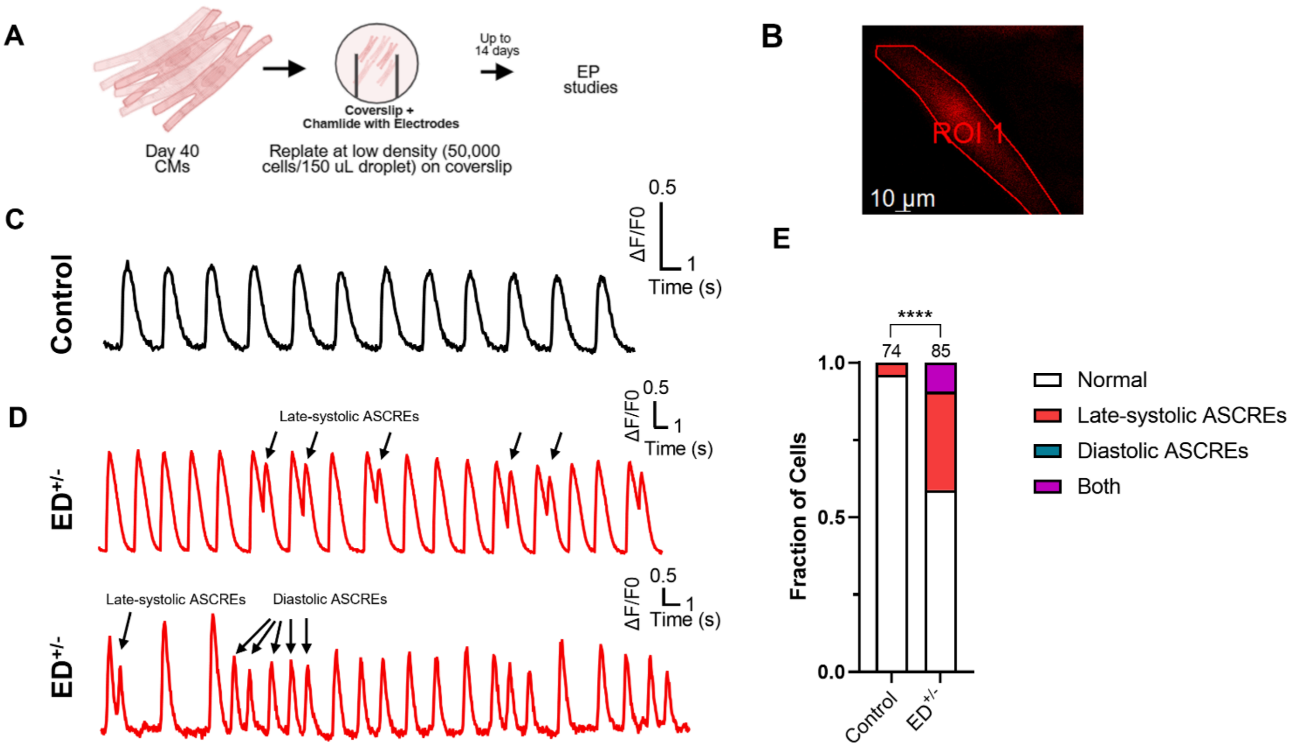


**Supplementary Figure 4.** Abnormal spontaneous $Ca^{2+}$ release events in individual E4146D$^{+/-}$ hiPSC-CMs**.** A) Workflow for low density hiPSC-CM replating for single cell or small cluster analyses. B) Sample $Ca^{2+}$ signal from an individual hiPSC-CM. C) Sample spontaneous $Ca^{2+}$ transient traces from a control hiPSC-CM. D) Sample spontaneous $Ca^{2+}$ transient traces from two separate E4146D$^{+/-}$ (ED$^{+/-}$) hiPSC-CMs, with one (top) showing late-systolic abnormal spontaneous $Ca^{2+}$ release events (ASCREs), and the other (bottom) showing both late-systolic and diastolic ASCREs. E) Fraction of control (n=74, 5 differentiation batches) and ED$^{+/-}$ (n=85, 6 differentiation batches) hiPSC-CMs showing ASCREs**.** Statistical comparisons were performed using the Chi-square test for categorical data.

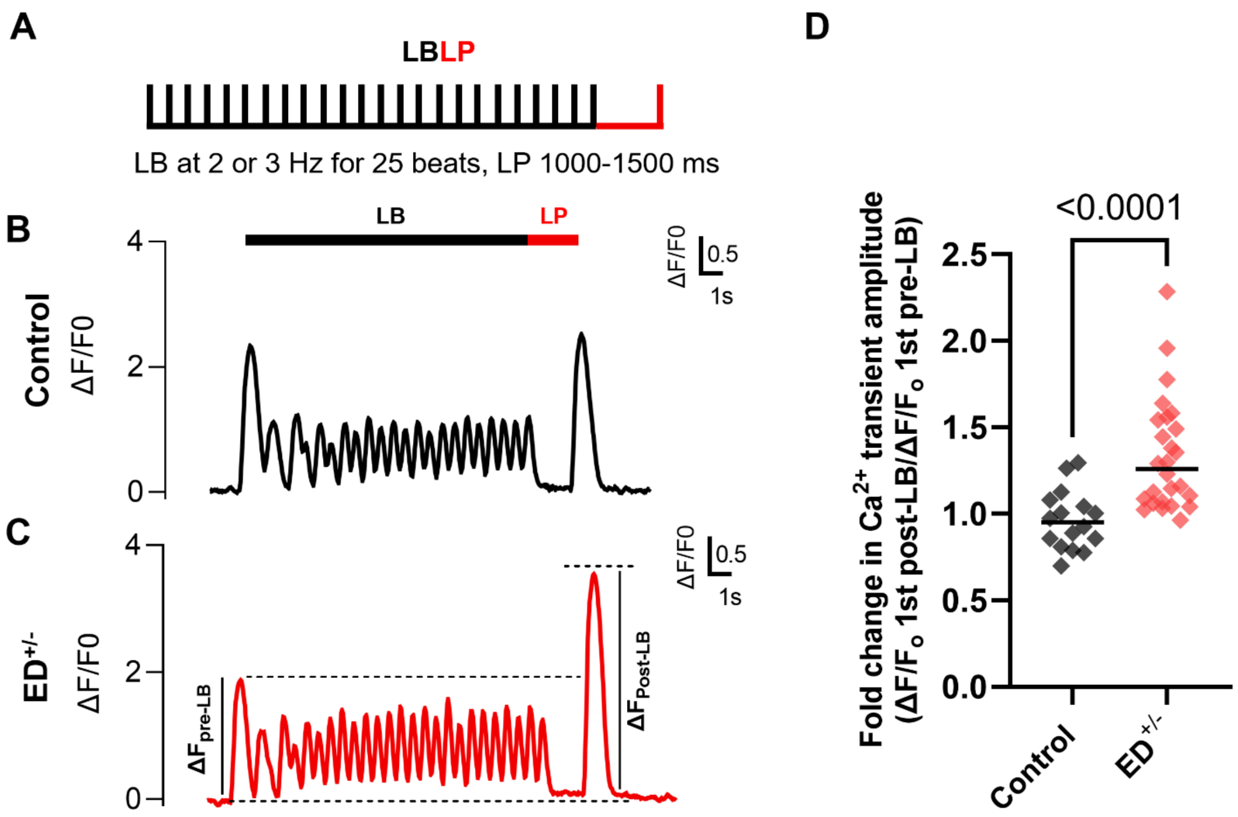


**Supplementary Figure 5.** Impact of long-burst pacing on individual control and E4146D$^{+/-}$ hiPSC-CMs. A) A long burst (LB) consisting of 25 consecutive, fixed-interval stimuli followed by a long pause (1000-1500 ms) ending with a stimulus (LP) was applied. Unlike in monolayers, which typically has a spontaneous beat with 1-3 seconds post-pacing, individual hiPSC-CMs typically had a longer post-pacing quiescence. Hence, we applied the LP stimulus to mimic a sinus beat shortly after a LB. B-C) Sample $Ca^{2+}$ transient traces in individual (B) control and (C) E4146D$^{+/-}$ (ED$^{+/-}$) hiPSC-CMs in response to the LBLP protocol, which shows a notably larger post-pacing $Ca^{2+}$ transient amplitude compared to pre-pacing. D) Quantification of the fold change of the post-LB $Ca^{2+}$ transient amplitude compared to the pre-pacing $Ca^{2+}$ transient amplitude in control (n=16, 4 differentiation batches) and ED$^{+/-}$ (n=26, 6 differentiation batches) hiPSC-CMs. Statistical comparisons were performed using the Kruskal-Wallis test for quantitative data.
